# Adjusting extracellular pH restores proteostasis and extends lifespan in a yeast model of polyglutamine toxicity

**DOI:** 10.1101/2025.09.07.674719

**Authors:** Khaleda Afrin Bari, Gabriela Nunes Marsiglio Librais, Martin L. Duennwald, Patrick Lajoie

**Author notes:** Corresponding author: Patrick Lajoie, PhD, Department of Anatomy and Cell Biology, The University of Western Ontario, London, Ontario, Canada N6A5C1.

## Abstract

Impaired proteostasis is a hallmark of aging and is associated with several neurodegenerative diseases, including Huntington’s Disease (HD) where the polyglutamine (polyQ) expanded Huntingtin aggregates to form insoluble inclusions bodies (IBs) associated with neurotoxicity. Chronological lifespan (CLS) in yeast resembles many aspects of aging of non-dividing cells such as neurons. During chronological aging, acidification of the culture media due accumulation of acetic acid is one of the major cell-extrinsic factors contributing to age-related cell death. Thus, buffering media pH to prevent acidification significantly extends longevity. Here, we found that cells expressing pathogenic polyQ expansion proteins display increased sensitivity to acetic acid and shortened CLS. Buffering media pH promotes both polyQ aggregation into IBs and promotes longevity. We also found that growth at alkaline pH induces the activation of heat shock response (HSR) in young cells. Such hormetic HSR activation subsequently allowed aged cells to mount a proper HSR in response to stresses such as heat shock or polyQ misfolding, leading to lifespan extension. Our study thus provides new insight into how pH can promote proteotoxic stress resistance and longevity by modulating the HSR.

## INTRODUCTION

There are at least 30 human disorders linked to age-associated accumulation of misfolded proteins including cancer, heart failure and many neurodegenerative diseases, such as Huntington’s Disease (HD) (Yang *et al*., 2014a; Meller and Shalgi, 2021a). Age-dependent accumulation of insoluble misfolded protein aggregates in humans (Yang *et al*., 2014b; Koyuncu *et al*., 2017; Martínez *et al*., 2017; Meller and Shalgi, 2021b) is a hallmark of several protein misfolding diseases and is recapitulated by many model organisms: yeast, worms, flies, and mice (Moronetti Mazzeo *et al*., 2012; Denoth Lippuner *et al*., 2014; Kumsta *et al*., 2017; Cabral-Miranda *et al*., 2022). Increased toxicity in protein misfolding diseases with age suggests a progressive deterioration of the protein homeostasis (proteostasis) network that regulates cellular protein quality control (Duennwald and Lindquist, 2008; Chafekar and Duennwald, 2012; Taylor and Dillin, 2013; Mesgarzadeh *et al*., 2022; Hipp and Hartl, 2024). Therefore, understanding the pathways that regulate protein folding and clearance of misfolded proteins is critical to develop new therapeutic strategies to improve our aging population’s healthspan.

HD (Huntington, 2003) is caused by an autosomal dominant mutation in the HTT gene on chromosome 4, which leads to an abnormally long CAG repeat sequence that produces a mutant huntingtin protein with an expanded polyglutamine tract (A novel gene containing a trinucleotide repeat that is expanded and unstable on Huntington’s disease chromosomes. The Huntington’s Disease Collaborative Research Group, 1993; Penney *et al*., 1997). This mutant protein forms toxic aggregates that primarily affect neurons in the striatum of the brain, disrupting cellular processes including transcription regulation, protein trafficking, and mitochondrial function (Gusella and MacDonald, 2000; Sawa, 2001; Fu *et al*., 2018). As the disease progresses, the abnormal protein accumulation leads to progressive neuronal dysfunction and eventually cell death (Hickey and Chesselet, 2003; Moujalled *et al*., 2021). The severity and age of onset correlate with the length of the CAG repeat expansion, with longer repeats typically resulting in earlier onset and more severe symptoms (Gusella and MacDonald, 2000; Roos, 2010).

Yeast is a powerful and well-established model to study aging, including basic aspects that are directly applicable to human aging (Kaeberlein *et al*., 2001; Chen *et al*., 2005; Fontana *et al*., 2010; Mirisola *et al*., 2014; Zimmermann *et al*., 2018). Yeast also allows the study of two genetically distinct paradigms of aging: chronological and replicative lifespan (Longo *et al*., 2012; He *et al*., 2018; Mirisola and Longo, 2022). Chronological lifespan in yeast is defined as the amount of time non-dividing cells survive in stationary phase, a paradigm relevant to the aging of post-mitotic cells such as neurons (Fabrizio and Longo, 2003; Laun *et al*., 2006; Longo and Fabrizio, 2012). During the course of chronological aging, yeast cells undergo distinct growth phases: the mid-log phase consists of a mixture of dividing cells and quiescent cells, diauxic shift (metabolic shift transitioning into stationary phase), and stationary phase (Mirisola and Longo, 2022). Each of these growth phases is characterized by distinct metabolic activities and gene expression profiles, which parallel central aspects of aging mammalian cells, such as increased extracellular pH, increased respiratory activity, arrested cell cycle, and the dysregulation of lipid and protein homeostasis (Mohammad *et al*., 2020). Indeed, we recently showed that activation of the unfolded protein response (UPR) to cope with endoplasmic reticulum stress is an important determinant of chronological lifespan (Chadwick *et al*., 2020a). Importantly, previously published studies show that HD-associated polyQ expansions shorten chronological aging (Cohen *et al*., 2016; Ruetenik *et al*., 2016; Pradhan *et al*., 2023).

The budding yeast model has proven very useful to determine the factors that control eukaryotic cell lifespan (Mirisola and Longo, 2022; Lin *et al*., 2025). Importantly, aging under standard culture conditions is linked to the pH of the culture medium. Under standard glucose culture conditions yeast cells accumulate fermentation products, such as ethanol, from sugars in aerobic culture (Fraenkel, 1981). Subsequent metabolism of ethanol results in increased media acidity through the secretion of organic acid. Initial pH in standard cultures goes from ∼5.8 to 3.0 once reaching stationary phase (Burtner *et al*., 2009a). Interestingly, while several other organic acids also accumulate in the culture medium during chronological aging, only acetic acid is sufficient to regulate chronological aging (Burtner *et al*., 2009b). When the extracellular pH is higher than 4.76, acetic acid is present mainly as acetate anions. Upon entering cells, acetic acid encounters the more alkaline cytoplasm, where it dissociates into acetate ions and protons. This process results in acetate accumulation and intracellular acidification, which can have multiple negative consequences. These effects include inhibiting cellular growth and metabolic processes like glucose consumption, as well as causing oxidative damage and depleting energy reserves within the cell and ultimately cell death (Ludovico *et al*., 2002; Fernández-Niño *et al*., 2015; Guaragnella and Bettiga, 2021). While the role of acetic acid in longevity has been extensively documented, the underlying mechanisms remain unclear. The interplay between pH, acetic acid and aging in yeast involves several mechanisms, including mitochondrial dysfunction, vacuolar acidification, modulation of nutrient-sensing pathways like TOR, and activation of stress response pathways (Burtner *et al*., 2009c; Yucel *et al*., 2014; Guerreiro *et al*., 2016; Dong *et al*., 2017; Chaves *et al*., 2021; Deprez *et al*., 2021). Understanding these mechanisms not only provides insights into yeast biology but also has potential applications in biotechnology and human health research, given the conservation of many of these pathways across species.

Many studies show that cellular stress responses are important modulators of polyQ toxicity (Lajoie and Snapp, 2011; Wu *et al*., 2014; Chadwick and Lajoie, 2019; Chadwick *et al*., 2020b; Das *et al*., 2023; Espina *et al*., 2023). Yet, the regulation of the HSR in aging cells is mostly unclear. The HSR is a conserved cellular response to various stresses, including exposure to elevated temperatures and the presence of misfolded proteins in the cytosol. In both yeast and humans, the HSR is primarily regulated by the highly conserved heat shock transcription factor, Hsf1 (Akerfelt *et al*., 2007; Dea and Pincus, 2024). During heat shock, Hsf1 translocates from the cytosol to the nucleus and binds as a homo-trimer to heat shock elements (HSEs), which are promoter sequences that regulate the expression of heat shock genes. The transcriptional activation of the HSR also involves Hsf1 phosphorylation, dephosphorylation, and acetylation, and interactions of Hsf1 with molecular chaperones, such as Hsp90 and Hsp70 (Morimoto *et al*., 1996; Pirkkala *et al*., 2001; Batista-Nascimento *et al*., 2011; Garde *et al*., 2024; Mühlhofer *et al*., 2024). Recently it was shown that orphan proteins activate Hsf1 by sequestering the chaperone Hsp70 and its co-regulator Sis1, which normally bind to and repress Hsf1 under non-stress conditions. When orphan proteins accumulate during stress, they titrate these chaperones away from Hsf1, thereby releasing it from repression and activating the heat shock response (Yanagitani *et al*., 2017; Krakowiak *et al*., 2018; Boos *et al*., 2019; Masser *et al*., 2019; Feder *et al*., 2021; Krämer *et al*., 2023; Ali *et al*., 2024). The HSR induces the expression of molecular chaperones, proteins of the ubiquitin proteasome system, and proteins involved in energy metabolism, to protect cells from stress. Several genetic studies indicate that the HSR plays a major role in healthy cellular aging and age-dependent impairment of the heat shock response is well documented in different organisms, including yeast, worms and flies and human cells, including human neurons (Morimoto *et al*., 1997; Hsu *et al*., 2003; Morrow *et al*., 2004; Kern *et al*., 2010; Ohtsuka *et al*., 2011; Watterson *et al*., 2022; Kovács *et al*., 2024). The accumulation of misfolded proteins, such as expanded polyQ proteins, may thus be enhanced by age-dependent defects in Hsf1 activation (Gomez-Paredes *et al*., 2021; Santiago and Morano, 2022; Kim and Gomez-Pastor, 2023; Tataridas-Pallas *et al*., 2024). Yet, on a cellular and genetic level, it remains unclear how aging impairs Hsf1 activation, thereby weakening the HSR, and ultimately increasing the toxicity of misfolded proteins. By using yeast genetics and chronological lifespan assays, we sought to determine the mechanism by which extracellular pH impacts protein quality control to regulate age-dependent polyQ aggregation and toxicity. Given the emergence of pH as an extrinsic modulator of cellular toxicity in the aging brain, our study identified new mechanisms that could contribute to polyQ toxicity across organisms.

## RESULTS

### Preventing media acidification extends lifespan of cells expressing expanded polyQ

Given that decreased extracellular pH due to accumulation of acetic acid is a major modulator of chronological aging in yeast (Burhans and Weinberger, 2009; Burtner *et al*., 2009a) (**Figure 1A**), we sought to determine if buffering culture media pH would impact longevity of cells expressing HD-associated expanded polyQ proteins. To do so, we took advantage of a well characterized yeast model of HD (Meriin *et al*., 2002, 2003; Duennwald *et al*., 2006b; Duennwald, 2013; Jiang *et al*., 2017; Albakri *et al*., 2018; Braun and Büttner, 2021) where the first exon of Htt (Htt^ex1^) fused to FLAG and CFP tags is regulated by *GAL1* (induced in presence of galactose) or *MET13* (induced in absence of methionine) promoters. The Htt^ex1^ sequence lacks the proline rich domain, which is a requirement to unveil polyQ toxicity in yeast (Duennwald *et al*., 2006b). First, we sought to determine if buffering the initial media pH would affect lifespan of cells expressing various lengths of polyQ proteins. In our design, we used 25Q as a non toxic and non disease-associated polyQ length that serves as the control for both *GAL1/MET13* expression systems. For the stronger *GAL1* expression system, we employed 46Q as our disease-associated allele. For the weaker *MET13* promoter, we use the 103Q. This approach has the advantage of using expanded polyQ with modest toxicity and growth defects in exponentially growing cells (Duennwald *et al*., 2006a; Jiang *et al*., 2017), allowing us to measure changes in toxicity during the aging process. We assessed chronological lifespan by culturing cells to stationary phase and measuring cell death over time using propidium iodide (Chadwick *et al*., 2016a, 2020c). To buffer the pH, we used two methods previously shown to extend lifespan in yeast: *i)* adjusting media pH with NaOH (Fabrizio *et al*., 2004) and *ii)* buffering using a low salt pH 6.0 MES buffer (Burtner *et al*., 2009a). Modulating extracellular pH did not affect growth and cells reached similar densities at day 3 (**Supplemental Figure 1**). Using these approaches we found that adjusting initial media to either 5.7 or 6.5 with NaOH significantly extended cell viability of the 46Q expressing cells during chronological aging (**Figure 1B and C**). Similarly, addition of MES also promotes longevity of the 46Q (**Figure 1D**). To ensure that the phenotypes observed are not dependent on the changes in carbon sources (from glucose to galactose), we assessed the effect of pH adjustment on aging using the *MET13* expression system. Similarly, we found that while 103Q displays a shorter lifespan than the 25Q counterpart, addition of MES extended longevity of both strains while eliminating the shorter lifespan phenotype associated with 103Q (**Figure 1E**).Finally, we assessed the ability of adjusting pH to extend lifespan using a regrowth assay on agar plate. We found that adjusting the pH of the aging culture to 6.5 also extended lifespan of the 46Q expressing cells when visualized by spotting cells on agar plate at different time points during the aging process (**Figure 1F**).

**Figure 1:**
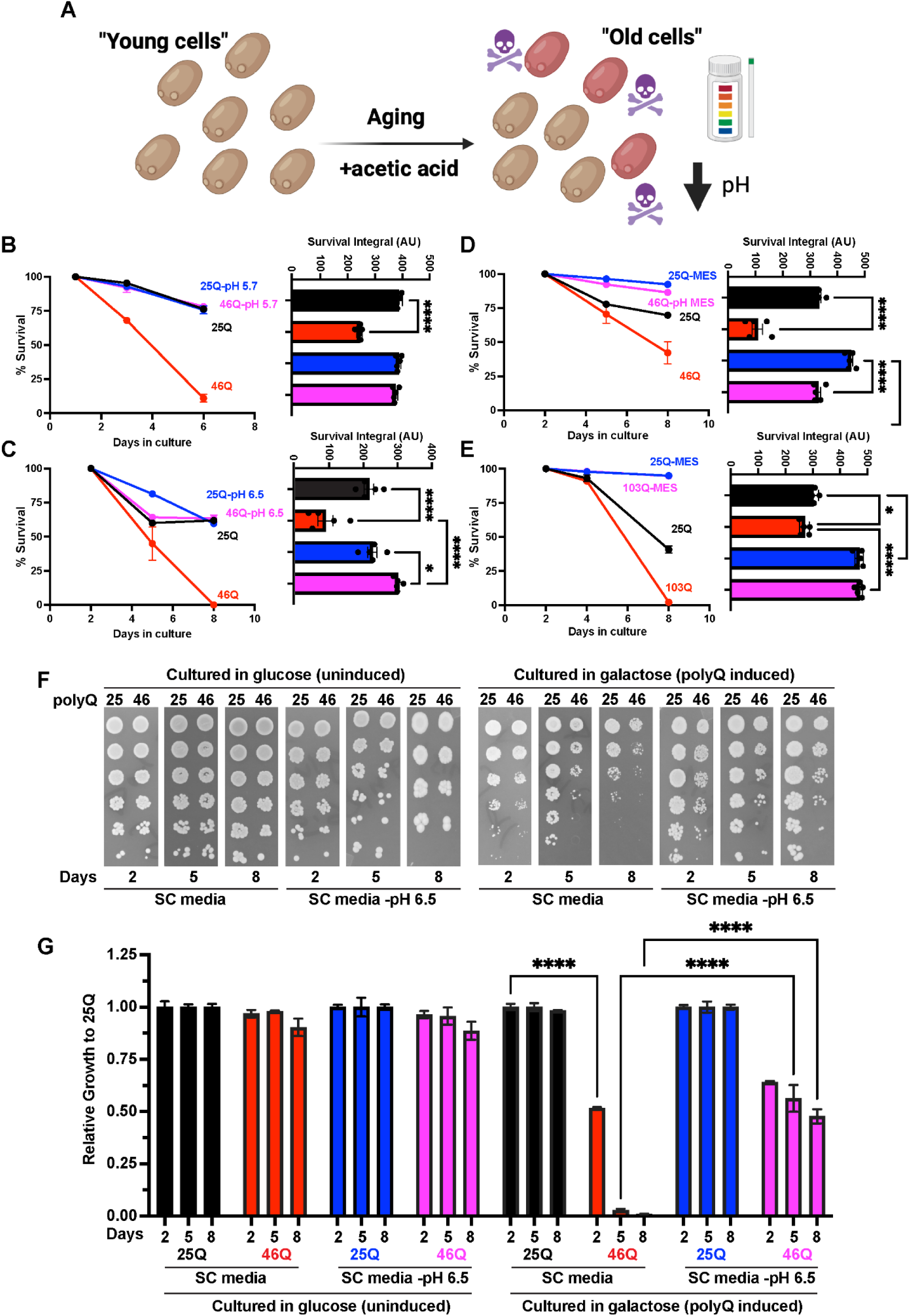
Buffering the pH of the aging culture extends the lifespan of cells expressing expanded polyQ. **A)** Chronological lifespan (CLS) is defined by the amount of time cells can survive at stationary phase. CLS is associated with rapid acidification of the culture medium, unmasking acetic acid toxicity. **B-D)** Cells expressing 25Q-CFP or either expanded 46Q-CFP were induced in galactose and processed for CLS in standard SC media or media buffered to pH 5.7 **(B)**, pH 6.5 **(C)** or buffered with MES pH 6.0 **(D)**. Cell viability was monitored over time using propidium iodide. Area under the curve (survival integral) is shown in bar graphs. **E)** Cells expressing 25Q-CFP or either expanded 103Q-CFP were induced in absence of methionine and processed for CLS in standard SC media or media buffered with MES pH 6.0. Cell viability was monitored over time using propidium iodide. **F)** Cells expressing 25Q-CFP or either expanded 46Q-CFP were induced in galactose and processed for CLS in standard SC media or media buffered to pH 6.0 and spotted on agar plates at different time intervals. **G)** Quantification of the aging assay on agar plates is shown in the bar graph. *p<0.05 ****p<0.0001 (two-way Anova followed by Tukey’s multi comparison test).

Overall, our data show that preventing media acidification extends the lifespan of cells expressing expanded polyQ. We reasoned that the shorter lifespan of the 46Q cells could be due to faster acidification of the media. First we measured the pH of aging cultures for both 25Q and 46Q over time. We found that pH decreased similarly over time for both strains (**Figure 2A**). Interestingly, when we performed the experiment using the longer and more toxic 72Q construct, we found that the pH did not decrease to the same extent, probably reflecting the rapid loss in viability associated with longer polyQ expansions (Jiang *et al*., 2017). We then reasoned that the shorter lifespan of 46Q cells could be due to increased sensitivity to acetic acid. Indeed, we found a correlation between polyQ length and sensitivity to acetic acid. Both 46Q and 72Q display reduced viability when challenged with acetic acid compared to 25Q (**Figure 2B**) suggesting decreased tolerance to acetic acid with the HD-associated polyQ expansions. Since polyQ toxicity is due to accumulation of toxic Htt^ex1^ (Lajoie and Snapp, 2010; Olshina *et al*., 2010; Sathasivam *et al*., 2010; Miller *et al*., 2011; Ormsby *et al*., 2013) oligomers, we next sought to assess whether changes in extracellular pH and subsequent toxic acetic acid accumulation have any impact on polyQ aggregation and formation of IBs.

**Figure 2:**
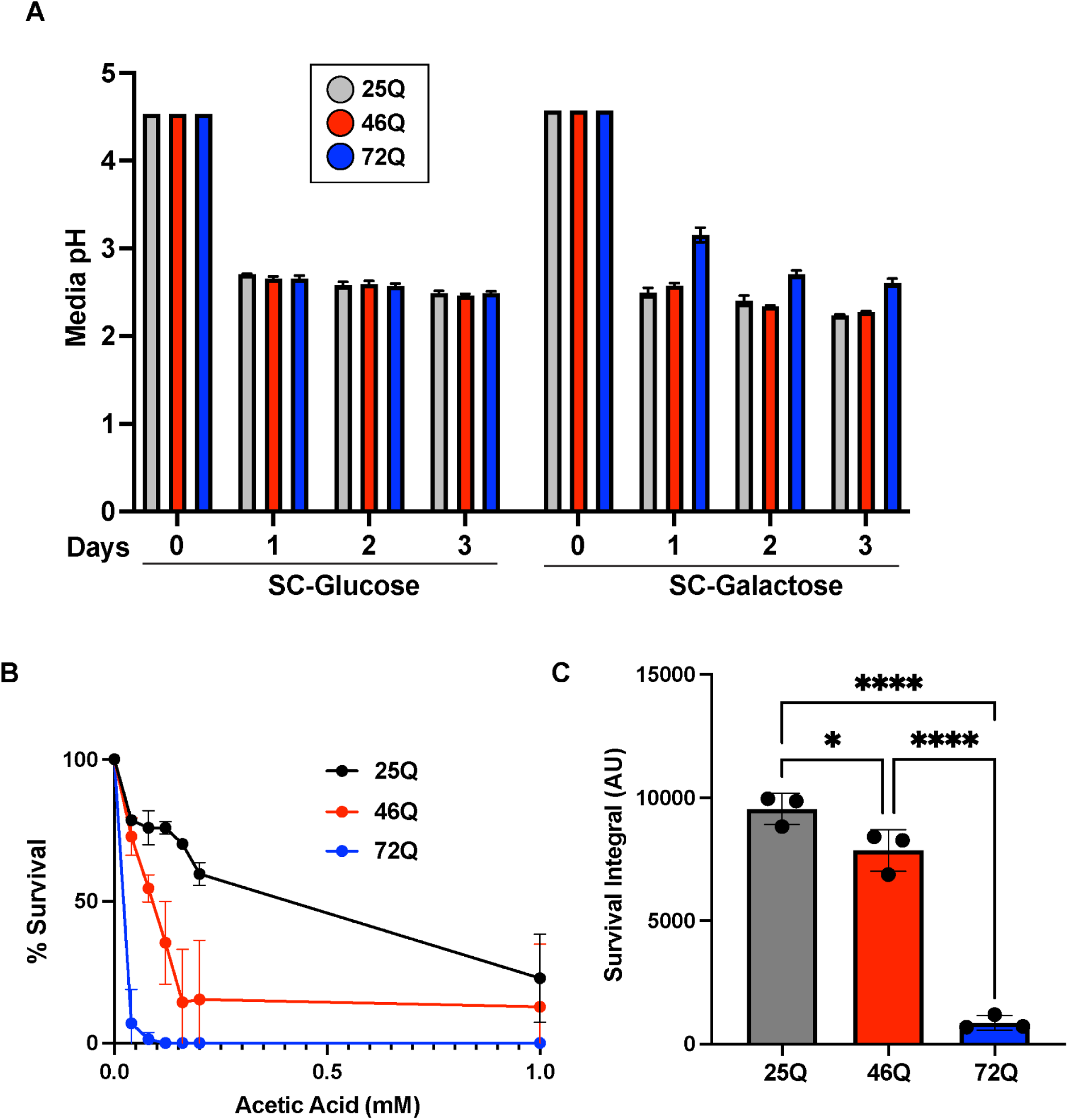
Cells expressing expanded polyQ display increased sensitivity to acetic acid. **A)** pH rapidly drops during chronological aging. Cells expressing 25Q-CFP or expanded 46Q-CFP and 72Q-CFP were aged in glucose (uninduced) or induced in galactose and processed for CLS in standard SC media. The pH of the cultures was monitored over time and shown in the bar graph (n=3 +/- SEM). **B)** Cells expressing 25Q-CFP or expanded 46Q-CFP and 72Q-CFP were aged for 3 days in galactose inducing media and treated with different concentrations of acetic acid. Viability was assessed with propidium iodide. Survival percentages normalized to untreated cells are shown on the graph (n=3 +/- SEM). The Area under the curve is shown in the bar graph (n=3 +/- SEM). *p<0.05 ****p<0.0001 (Anova followed by Tukey’s multi comparison test).

### Extracellular pH modulates polyQ aggregation

In yeast, several factors can impact the ability of polyQ proteins to oligomerize and aggregate into IBs. For example, the presence of the proline rich region of Htt^ex1^ leads to formation of compact and benign IBs while absence of the proline rich region is associated with morphologically distinct amorphous IBs associated with the toxic phenotype (Duennwald *et al*., 2006b). PolyQ aggregation and toxicity in yeast is also dependent on the Rnq1 prion (Meriin *et al*., 2002; Duennwald *et al*., 2006a; Gropp *et al*., 2022). Meanwhile, polyQ aggregation status has been shown to be altered during chronological aging (Cohen *et al*., 2012). Thus, we sought to determine the impact of extracellular pH on polyQ aggregation during the aging process. To do so, we imaged both 25Q and 46Q tagged with CFP over the course of aging in both regular media and in media buffered with MES. While 25Q remains diffuse in the cytoplasm, we found that 46Q assembles into IBs. Surprisingly, cells aged in buffered media displayed increased aggregation at day 3 (**Figure 3A and B**). We found similar results in cells aged in media buffered to pH 6.5 with NaOH (**Supplemental Figure 2**). Interestingly, 46Q cells grown in unbuffered media display a decreased proportion of cells with IBs at day 3 (**Figure 3B**). Given the ability of buffered media to reduce the gradual decline in cell viability during the course of aging, these results indicate a protective role for IBs formation during chronological lifespan. These results are consistent with the findings showing that intermediate soluble oligomers rather than IBs represent the toxic species in HD (Behrends *et al*., 2006; Takahashi *et al*., 2008; Lajoie and Snapp, 2010; Miller *et al*., 2011; Leitman *et al*., 2013; Kim *et al*., 2016; Gropp *et al*., 2022; Miguez *et al*., 2023).

**Figure 3:**
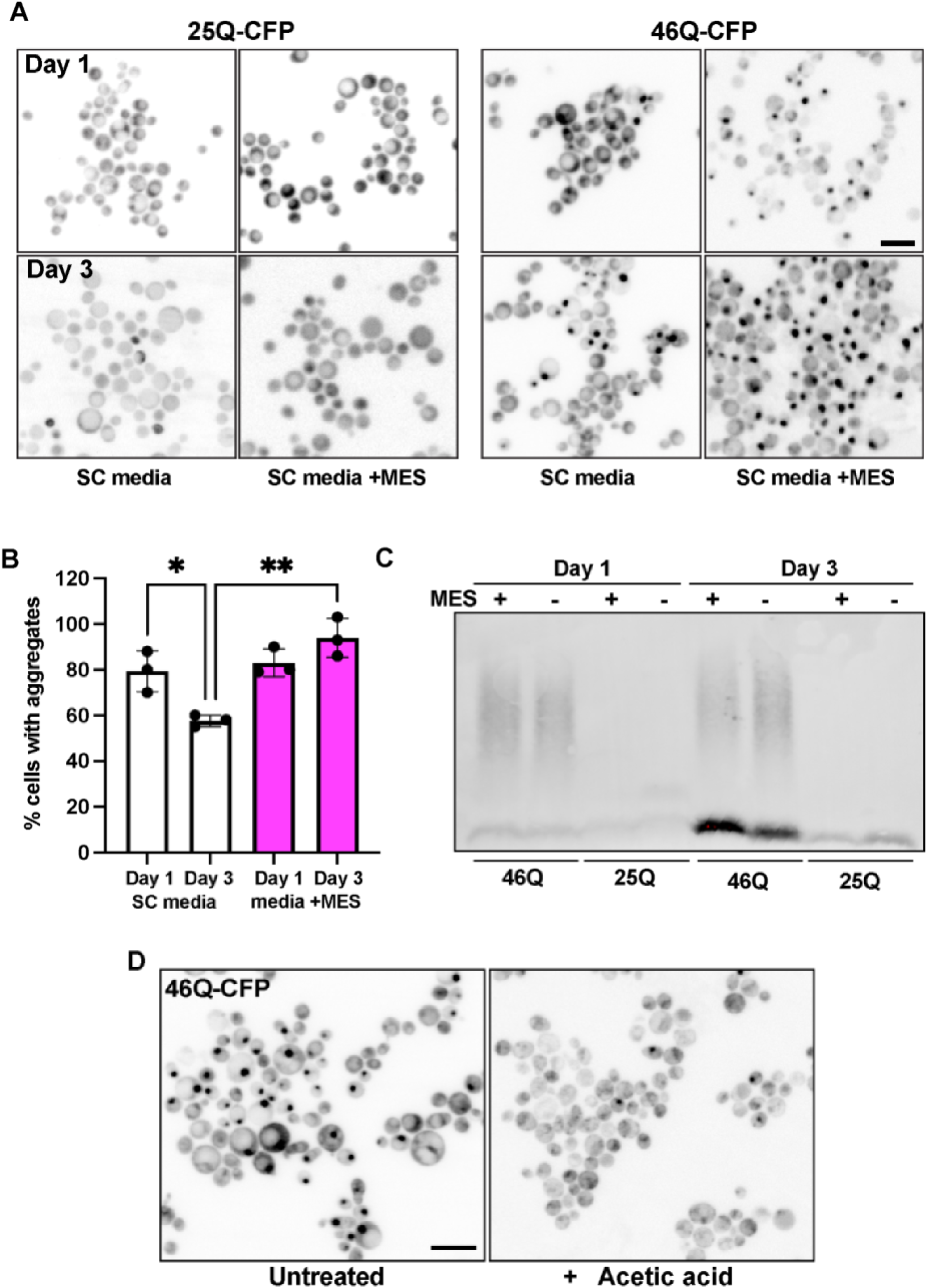
Buffering the pH of aging cultures increases polyQ IBs formation. **A)** Cells expressing 25Q-CFP or either expanded 46Q-CFP were processed for CLS in standard SC media or media buffered with MES containing galactose. Fluorescence images were collected on day 1 and 3. **B)** The percentage of 46Q-CFP expressing cells presenting visible IBs is presented in the bar graph (n=3). **C)** Cells expressing 25Q-CFP or either expanded 46Q-CFP were processed for CLS in standard SC media or media buffered with MES containing galactose. Cells were lysed on day 1 and 3 and processed for SDD-AGE. D) Cells expressing 46Q-GFP were induced overnight in SC media containing galactose and treated with 0.06 mM acetic acid and images collected after 90 min of treatment. Bars: 10µm. *p<0.05 **p<0.01 (Anova followed by Tukey’s multi comparison test).

### Growth at higher pH remodels the proteostasis network

Given our results showing that growth buffered media alters polyQ toxicity and aggregation, we next sought to determine the effect of growing cells at higher pH on the transcriptome. First, we cultured wild-type yeast cells at both regular SC media and SC media buffered to pH 6.5 with NaOH. Total RNA was isolated from cells after 16 hours of growth (day 1) as the cells are transitioning into the post-diauxic shift and changes in the transcriptome assessed by RNA-seq. From these dataset we identified 291 downregulated and 284 upregulated genes (p<0.05) in cells grown at pH 6.5 when compared to normal growth conditions (**Figure 4A and Supplemental File 1**). Cells cultured at different pH show distinct transcriptional signatures as seen by the principal component analysis (**Figure 4B**). Gene ontology (GO) analysis of upregulated genes revealed enrichment in genes involved in ribosome biogenesis (**Figure 4C**). Repression of ribosome biogenesis is linked to glucose exhaustion (Yin *et al*., 2003; Lee and Lee, 2008) in the growth media and is a signature of cells entering the stationary phase (Zhang *et al*., 2023) and proteotoxic stress (Li *et al*., 2000; Guerra-Moreno *et al*., 2015; Albert *et al*., 2019). Similar analysis of downregulated genes revealed reduced expression of genes associated with protein folding and expression of heat shock proteins (**Figure 4D**). Since repression of ribosome biogenesis and induction of the heat shock response (HSR) are both indicators of cellular stress, we reason that cells grown at lower pH are protected against stress associated with chronological aging. Moreover, we found that extracellular pH still significantly decreases during aging, even in buffered media (**Supplemental Figure 3**). This suggests that young cells grown at higher pH may experience long lasting changes that could be beneficial for promoting longevity. We also assessed the effect of growth in buffered media on the intracellular cytosolic pH using the genetically encoded reporter pHLurorin (Miesenböck *et al*., 1998; Reifenrath and Boles, 2018). We found that buffering media pH to 6.5 did not significantly impact intracellular pH (**Supplemental Figure 4**) suggesting the effect of changes in pH on the HSR may be traced back to the cell surface/cell wall.

**Figure 4:**
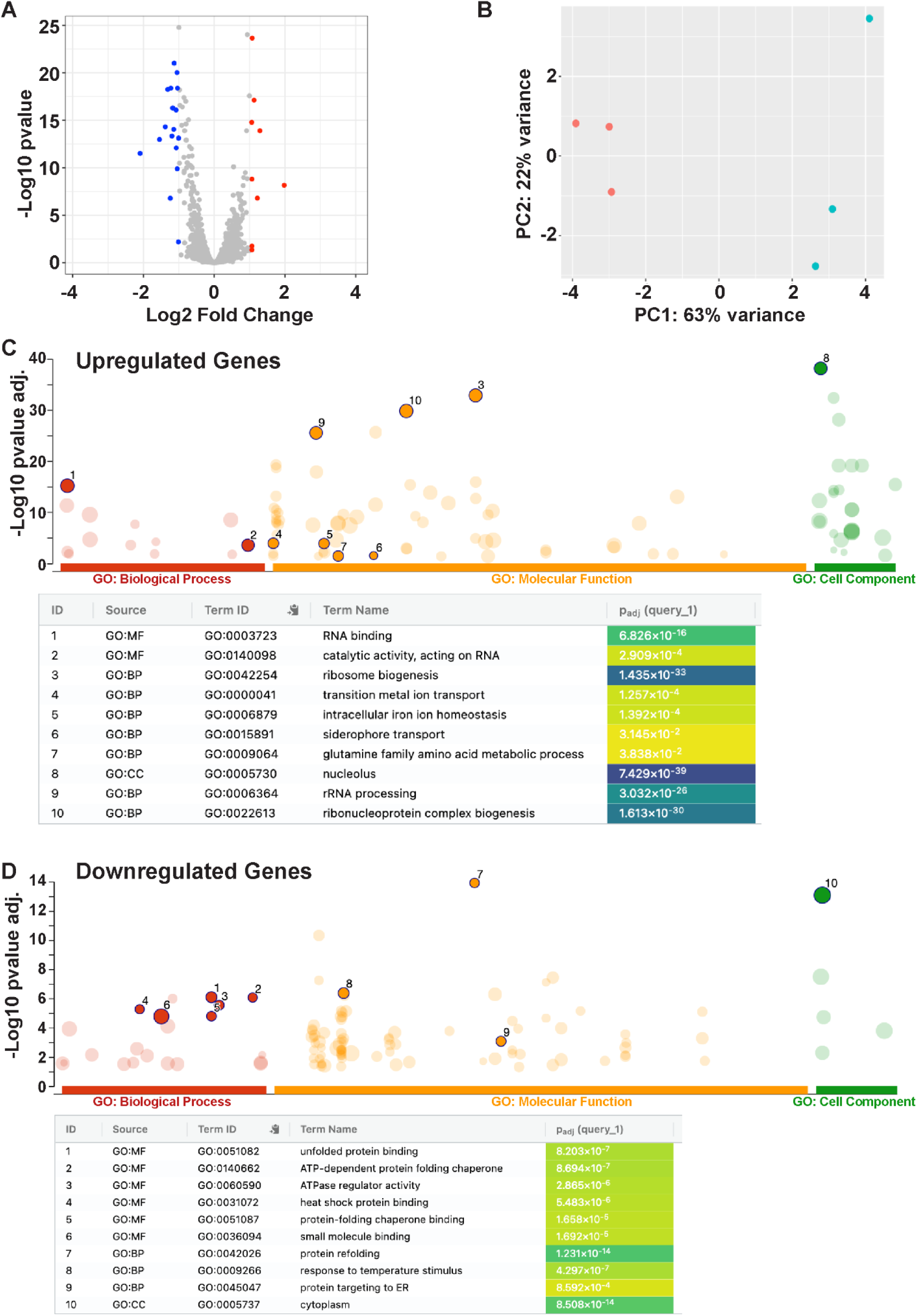
Growth in buffered media reduced expression of genes associated with proteotoxic stress at day 1. **A)** Volcano plot of changes in gene expression as determined by RNA-seq in W303 yeast cells grown for 16 hours in either standard SC media or SC media buffered to pH 6.5. Differentially expressed genes with an adjusted *P*-value <0.05 and log2 fold change > 1.0 are colored red (up regulated in buffered media relative to standard SC) and blue (down regulated in buffered media relative to standard SC). **B)** Principal component analysis of centered log ratio normalized reads from cells grown in either standard SC media or SC media buffered to pH 6.5. Each point represents one biological replicate (*n* = 3). **C)** Manhattan plot illustrating the GO term enrichment analysis from the set of significantly upregulated genes with adjusted *P* < 0.05 in cells grown for 16 hours in either standard SC media or SC media buffered to pH 6.5.The top 10 GO terms are presented in the table. **D)** Manhattan plot illustrating the GO term enrichment analysis from the set of significantly downregulated genes with adjusted *P* < 0.05 in cells grown for 16 hours in either standard SC media or SC media buffered to pH 6.5.The top 10 GO terms are presented in the table.

One possibility is that cells grown at higher pH experience a preadaptive or hormetic stress that increases their resistance along the aging process. To investigate this possibility, we followed the induction of the HSR after cells are inoculated in either standard SC media or media at buffered pH 6.5 (referred here as Day 0). Induction of the HSR can be monitored in living cells using a yeast strain expressing a yellow fluorescent protein (YFP) under the control of a promoter containing 4 tandem heat shock response elements (4xHSE). Mean fluorescence intensity is monitored over time using flow cytometry. Using this tool, we found that indeed, cells grown at pH 6.5 display rapid induction of the HSR compared to cells in standard media, which plateaued at 6h post inoculation (**Figure 5A**). As a control, no significant change in fluorescence intensity was observed for a strain expressing GFP from a constitutive GPD promoter (**Figure 5B**). In support of induction of proteotoxic stress we also observed increased aggregation of the disaggregase Hsp104 (Shorter, 2008; Mack *et al*., 2023) when cells were inoculated in buffered media (**Supplemental Figure 4**) reflecting increased accumulation of misfolded substrates. We postulate that HSR activation in young cells protects them against future proteostatic insults along the aging process. To test this hypothesis, we exposed cells grown in normal media or media at pH 6.5 to heat shock at 42°C at day 1 and 3 of the aging process. On day 1, both cell populations displayed increased HSR activation upon heat shock (**Figure 5C**). While cells grown in normal media lose the ability to induce the HSR at day 3, cells grown at pH 6.5 remain competent to induce the HSR (**Figure 5C**). Notably, unlike the HSR, media pH buffering had only a modest effect on induction of the general stress response (GSR) and no significant changes were observed for the response to endoplasmic reticulum stress (through the unfolded protein response) (**Supplemental Figure 5**). Taken together, our data suggest that preventing media acidification leads to remodeling of the proteostasis network (with a preference for the HRS) and preserves the ability of aged cells to respond to proteotoxic stress. Given the dysfunctional HSR is a hallmark of HD, we next sought to determine how polyQ expansions and acid stress impact the HSR during lifespan.

**Figure 5:**
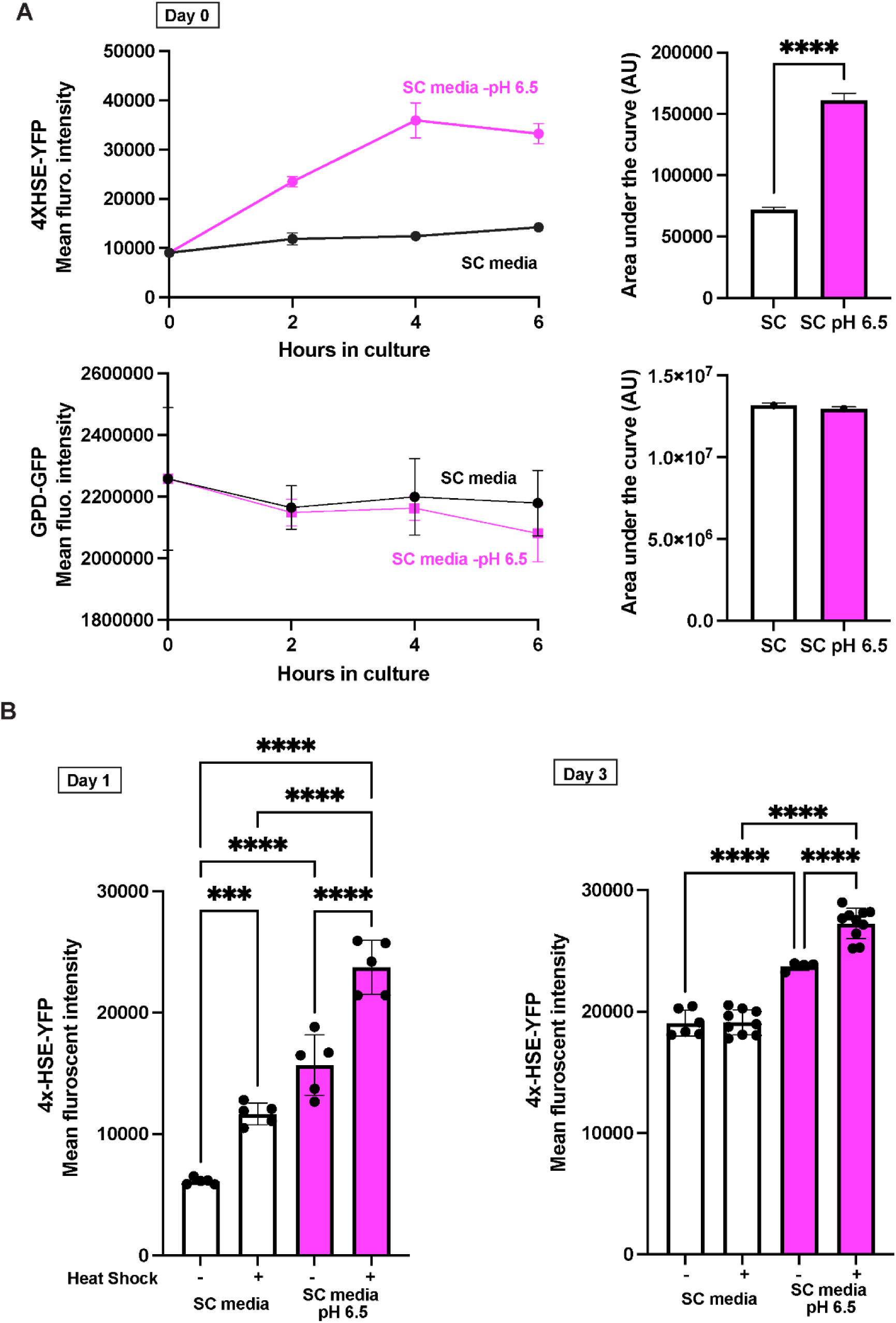
Growth at higher pH activates a protective HSR. **A)** Fresh culture of cells expressing either a HSR responsive 4xHSE-YFP reporter (Top) or the constitutively expressed GFP under the GPD promoter (bottom) were grown overnight and transferred to either standard SC media or SC media buffered to pH 6.5 (Day 0) and analysed by flow cytometry at different time intervals. Mean fluorescence intensity of each reporter is shown in the graph (n=5). Area under the curve is shown in bar graphs. B) Aging in buffered media preserved the ability of old cells to activate the HSR. Cells expressing the HSR responsive 4xHSE-YFP reporter were aged for 1 or 3 days followed by heat shock at 42°C for 60 min and processed for flow cytometry. Mean fluorescence intensity of each reporter is shown in the bar graph (n=3). ***p<0.005 ****p<0.001. (Anova followed by Tukey’s multi comparison test).

### Growth at higher pH improved stress tolerance of cells expressing expanded polyQ

First, we explored how young cells respond to pathological polyQ expansion. We used RNA-seq to reveal the effects of expanded polyQ on cells at day 1. In this data set we found 91 downregulated and 9 upregulated genes (p<0.05 and log_2_ fold change > 1.0) (**Figure 6A and Supplemental File 2**). Promoter analysis of downregulated genes revealed enrichment in stress responsive genes controlled by Hsf1 and Msn2/4 that control induction of the HSR and the general stress response respectively (**Figure 6B**). No enrichment was found for upregulated genes. GO term analysis of downregulated genes revealed enrichment for terms associated with chronological aging and quiescence such as glycogen and trehalose metabolism (Enjalbert *et al*., 2000; Shi *et al*., 2010; Leonov *et al*., 2017; Mohammad *et al*., 2020) (**Figure 6C**). Specifically, we found downregulation of genes such as *TSL1* and *TSP2* which encode subunits of the trehalose 6-phosphate synthase complex (De Virgilio *et al*., 1993; Vuorio *et al*., 1993), *GLC3*, a glycogen branching enzyme (Rowen *et al*., 1992) involved in glycogen accumulation during aging and *GSY1* (Farkas *et al*., 1990), which encode glycogen synthase (**Figure 6D**). Consistent with the defect in HSR activation associated with polyQ expansions (Hay *et al*., 2004; Labbadia *et al*., 2011; Chafekar and Duennwald, 2012; Riva *et al*., 2012; Gomez-Pastor *et al*., 2017; Kim and Gomez-Pastor, 2023; Mansky *et al*., 2023), we also found downregulation of several genes encoding heat shock proteins (*HSP104, HSP42, HSP26*) (**Figure 6E**).

**Figure 6:**
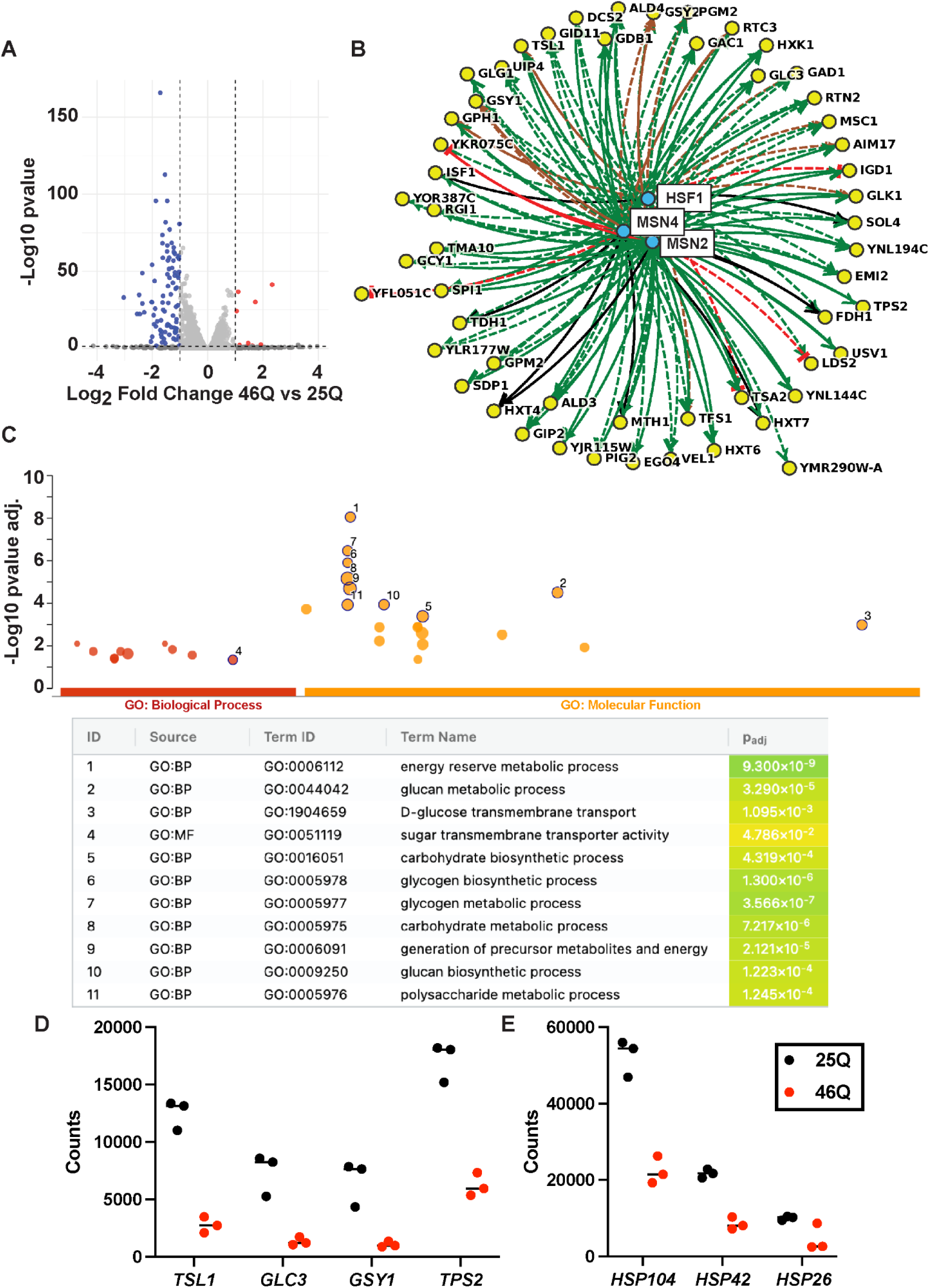
Cells expressing expanded polyQ display reduced expression of genes associated with the HSR. **A)** Volcano plot of changes in gene expression as determined by RNA-seq in yeast cells expressing either 25Q or 46Q-CFP grown for 16 hours (day 1). Differentially expressed genes with an adjusted *P*-value <0.05 and log2 fold change > 1.0 are colored red (up regulated in 46Q relative to 25Q) and blue (down regulated in 46Q relative to 25Q). **B)** Differentially downregulated genes in 46Q were analysed for transcription factor associations using YEASTRACT. Hsf1 and Msn2/4 associations with downregulated genes in the 46Q strain. The experimental evidence underlying each regulatory association (full lines for DNA-binding evidence; dashed for expression evidence), as well as the sign of the interaction—positive (green), negative (red), positive and negative (brown), or undefined (black) are displayed. **C)** Manhattan plot illustrating the GO term enrichment analysis from the set of significantly downregulated genes with adjusted *P* < 0.05 and log_2_ fold > 1.0 in 46Q-CFP expressing cells cells.The top 11 GO terms are presented in the table. **D)** Normalized RNA sequencing read counts are shown for *TSL1*, *GLC3*, *GSY1* and *TPS2* in cells expressing 25Q and 46Q-CFP. **E)** Normalized RNA sequencing read counts are shown for *HSP104*, *HSP26* and *HSP42* in cells expressing 25Q and 46Q-CFP.

Given these results, we next asked whether buffering the culture media to pH 6.5 would affect the ability of polyQ expressing cells to induce the HSR. Indeed we found that old cells (day 3) expressing 46Q display significant increases in both *HSP104* and *HSP42* levels when cultured at pH 6.5 (**Figure 7A and B**). To confirm that the HSR is required to alleviate polyQ toxicity in yeast, we then expressed both 25 and 46Q constructs in a strain carrying a truncated *HSF1* allele lacking the C-terminal domain (*HSF1 ΔC*) that prevents growth at high temperature (Morano *et al*., 1999; Truman *et al*., 2007). The HSF1 *HSF1 ΔC* strain displays reduced tolerance to acetic acid (**Supplemental Figure 6**), suggesting the HSR must play a role in the response to media acidification. When expressed in this strain, low expression of 103Q under the *MET13* promoter shows reduced growth compared to the *HSF1* wild-type counterpart (**Figure 7C and D**). Interestingly, growth at high pH did not rescue growth, suggesting that the ability of buffered media to alleviate polyQ toxicity partially depends on Hsf1 function. Supporting an impaired HSR in expanded polyQ expressing cells, we found that while showing no growth defect at 30°C, cells expressing 103Q display severe growth defect following heat shock at 42°C (**Figure 7 E and F**). Consequent with the ability of reduced polyQ toxicity and restored HSR activation at pH 6.5, we observed improved growth of 103Q expressing cells following heat shock in buffered media compared to standard culture conditions.

**Figure 7:**
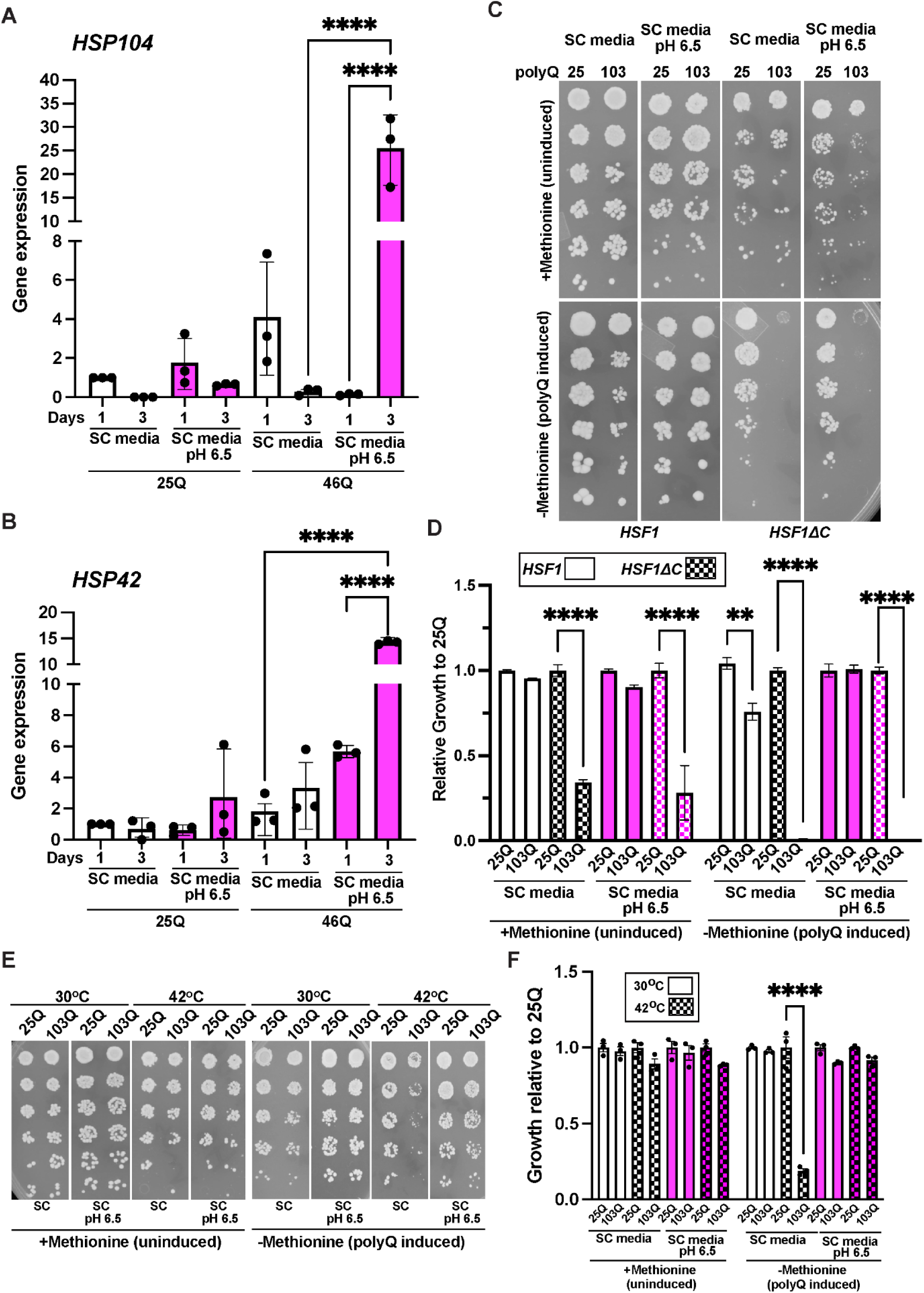
The HSR is required for protection against proteotoxic stress at higher pH. A and. **B)** Buffering media to pH 6.5 is associated with increased expression of heat shock proteins in 46Q cells at day 3. Cells expressing either 25Q-CFP or 103Q-CFP were processed for CLS and expression of *HSP104* **(A)** and *HSP42* **(B)** was assessed at day 1 and 3 using qRT-PCR. **C)** Wild Type *HSF1* or *HSF1ΔC* cells expressing either 25Q-CFP or 103Q-CFP were spotted on SC media or SC media buffered to pH 6.5 agar plates with (polyQ induced) or without (uninduced) methionine. **D)** Growth relative to uninduced 25Q on regular SC media is shown in the bar graph (n=3 +/-SEM). **E)** Cells expressing either 25Q-CFP or 103Q-CFP were induced for 16 hours with (polyQ induced) or without (uninduced) methionine in SC media or SC media buffered to pH 6.5. Subsequently cells were subjected to heat shock at 42°C for 1 hour and plated on regular SC media plates with or without methionine. **F)** Growth relative to uninduced 25Q on regular SC media is shown in the bar graph (n=3 +/-SEM). ***p<0.005 ****p<0.001. (Anova followed by Tukey’s multi comparison test).

Our findings demonstrate that buffering media pH to prevent acidification significantly extends the lifespan of cells expressing expanded polyQ proteins, while promoting the formation of protective inclusion bodies. Additionally, we showed that growth at higher pH induces hormetic activation of the heat shock response in young cells, which subsequently enables aged cells to mount an appropriate stress response to challenges such as heat shock or polyQ misfolding, ultimately leading to extended longevity and reduced proteotoxicity.

## DISCUSSION

### pH regulation and stress responses in chronologically aged cells

Yeast is an established model to determine the molecular mechanisms underlying the aging process (Mirisola *et al*., 2014; Dahiya *et al*., 2020). Work in yeast has enabled researchers to make several key discoveries regarding cellular mechanisms that regulate longevity (Mirisola *et al*., 2014; Wierman and Smith, 2014; Eleutherio *et al*., 2018). During chronological aging, rapid media acidification due to accumulation of acetic acid results in cellular toxicity. While other organic acids accumulate in the media of chronologically aged cultures, it was shown that only acetic acid is associated with toxicity (Burhans and Weinberger, 2009; Burtner *et al*., 2009a; Murakami *et al*., 2011). The fact that the extracellular pH rapidly drops even after inoculation in buffered media (**Supplemental Figure 1B**) suggests that exposure to higher pH early during the aging process can have long lasting effects to promote longevity. However, it was shown that buffering the pH of the aging culture, even several days after the original inoculation is sufficient to extend lifespan (Burtner *et al*., 2009a). In light of our results, one could think that such manipulation could still rewire the proteostasis networks to alleviate acid stress. It is attractive to think that lifespan extension could be a combination of both hormetic adaptation and acute prevention of acid stress.

Cells at stationary phase have been shown to accumulate heat shock proteins such as Ssa4 (Chughtai *et al*., 2001; Wasko *et al*., 2013) and Hsp90 co-chaperones (Tapia and Morano, 2010). Similarly, expression of small heat shock proteins such as Hsp26 and Hsp42 increase at stationary phase (Gasch *et al*., 2000) and Hsp42 accumulates in stress granule where they contribute to spatially partition proteins such as the histone deacetylase Hos2 during growth at stationary phase (Liu *et al*., 2012). Together with our data showing increased acetic acid sensitivity in the *HSF1 ΔC* strain (**Supplemental Figure 5**) we postulate that proteotoxic stress response is indeed required for proper chronological aging and manipulating extracellular pH can help rewire the protein quality control network to adapt cells to stress associated with aging. However, it was shown that buffering media to pH 9.0 results in shorter lifespan, suggesting an optimal threshold for the promotion of longevity exists (Murakami *et al*., 2011). Of note, buffering the pH of the extracellular media does not extend replicative lifespan, suggesting this phenomenon is rather restricted to non-dividing cells (Wasko *et al*., 2013).

pH regulation is closely linked to induction of the HSR and a transient acidification of the cytoplasm is required for activation of Hsf1 under heat stress (Weitzel *et al*., 1987; Triandafillou *et al*., 2020). Similarly, acid stress has been linked to activation of the HSR and thermotolerance (Coote *et al*., 1991; Cheng and Piper, 1994; Samanfar *et al*., 2017; Salas-Navarrete *et al*., 2023). Hsf1 trimerization, a key step in the activation of the HSR, is also pH dependent (Zhong *et al*., 1999; Choi *et al*., 2024). Conversely, alkalinization of the media also induces stress responses such as the ESR (Serrano *et al*., 2002; Ariño, 2010; Casamayor *et al*., 2012). How modulation of the extracellular pH during chronological aging affects intracellular pH remains elusive. Previous studies have linked changes in intracellular pH with cell growth regulation. For example, intracellular pH is tightly regulated and drops rapidly after glucose exhaustion (Dechant *et al*., 2010; Orij *et al*., 2012; Gutierrez *et al*., 2022) or energy depletion (Munder *et al*., 2016); acting as an integrative signal to dictate cell fate. Another example is the ability of intracellular pH to trigger lipid synthesis through the protonation of phosphatidic acid to couple membrane biogenesis to nutrient availability (Young *et al*., 2010). Given that we did not observe changes in intracellular pH in cells grown in buffered media, our results suggest that activation of the HSR at higher pH might be traced back to the cell surface or the cell wall. Indeed, cell wall integrity, stress tolerance and the heat shock response are closely associated (Imazu and Sakurai, 2005; Truman *et al*., 2007; Ribeiro *et al*., 2022). Acid stress is known to affect cell wall structure and composition (Cabib *et al*., 1989; Kapteyn *et al*., 2001). Relevant to our study, raising the pH from 4.0 to 6.0 reduces the relative proportion of β1,6-glucans within the β-glucan fraction of the cell wall, resulting in enhanced susceptibility to zymolyase digestion in cells cultured at pH 6.0 (Aguilar-Uscanga and François, 2003). Thus, one could postulate that changes in cell wall dynamics/composition could account for some of the phenotypes observed here.

Finally, what happens during chronological aging is complicated by the heterogeneous nature of the aging culture which contains both quiescent and non-quiescent cells with very different characteristics (Allen *et al*., 2006; Davidson *et al*., 2011; Sagot and Laporte, 2019; Mohammad *et al*., 2020). More work will be required to determine how cell extrinsic factors such as extracellular pH can affect both quiescent and non-quiescent cells within the aging yeast population.

### PolyQ expansions and aging

Here, we found that the extended longevity due to buffering of the media pH was associated with increased presence of polyQ IBs (**Figure 4**). These results support a protective role for IBs and the toxic nature of intermediate oligomers across HD models (Behrends *et al*., 2006; Takahashi *et al*., 2008; Lajoie and Snapp, 2010; Miller *et al*., 2011; Leitman *et al*., 2013; Kim *et al*., 2016; Gropp *et al*., 2022). Loss of proteostasis is a hallmark of age-dependent protein misfolding diseases, including HD (Douglas and Dillin, 2010; Klaips *et al*., 2018). This loss of proteostasis, can in turn modulate the ability of cells to incorporate polyQ proteins into IBs, leading to the appearance of toxic oligomers (Gidalevitz *et al*., 2006). Thus, by promoting proteostasis, buffering media pH appears to switch the balance back to protective IBs, extending lifespan. PolyQ aggregates are known to sequester a vast number of proteins leading to significant cellular dysfunction (Hosp *et al*., 2017). In the future, efforts should be made to better define the nature and compositions of the different polyQ IBs in yeast (Gruber *et al*., 2018) and how these relate to longevity. Interestingly, it was shown that the levels of the Hsp70 co-chaperone Sis1 regulates the nature of polyQ IBs. When present at elevated levels, Sis1 prevents the formation of dense IBs, directing them instead into permeable condensates where Hsp70 can accumulate in a liquid-like state, resulting in HSR activation (Klaips *et al*., 2020). How this mechanism impacts aging remains to be determined.

Similarly, how gene deletion known to extend or shorten lifespan such as *Δsch9* and *Δavr1* among others or other cell non-autonomous factors such as nutrients and sugar levels (Fabrizio *et al*., 2001; Powers *et al*., 2006; Matecic *et al*., 2010; Garay *et al*., 2014; Smith *et al*., 2016; Enriquez-Hesles *et al*., 2021; Schmiedhofer *et al*., 2025), impact polyQ aggregation and age-dependent toxicity should be better defined. Finally, caloric restriction has been shown to reduce the accumulation of acetic acid and shift the cell metabolism from fermentation to respiration (Ocampo *et al*., 2012). Therefore, it is reasonable to postulate that reducing acetic acid toxicity, either by neutralizing its effect by reducing acidification or lowering its production by moving away from fermentation, allows for lifespan extension.

### pH as a modulator of polyQ toxicity in mammals

Maintenance of neuronal intracellular pH is a tightly regulated process that involves multiple proteins and organelles (Brookes, 1997; Chesler, 2003). Metabolites such as lactate are essential for brain metabolism and regulates signaling and communication between neurons and astrocytes (Tang *et al*., 2014; Magistretti and Allaman, 2018). Lactate is metabolized by lactate dehydrogenase, which converts lactate to pyruvate and its activity is critical for aging and memory (Ross *et al*., 2010; Long *et al*., 2020; Frame *et al*., 2023). Both lactate and pyruvate can change intracellular pH and are metabolized inside cells. Lactate metabolism has been proposed to play a key role in several neurodegenerative diseases including Alzheimer’s, Parkinson’s and Huntington’s (Ross *et al*., 2010; Majdi *et al*., 2016; Al Ojaimi *et al*., 2024; Yang *et al*., 2024). Reduction in pH in the brain has also been linked with aging. A reduction of pH in vivo has been observed in older healthy patients with magnetic resonance spectroscopic imaging (Forester *et al*., 2010). Recently, it was shown that pH decreases in the brain of both patients and rodent models of Alzheimer’s disease (Decker *et al*., 2021). Interestingly, patients with Alzheimer’s disease have a significant increase in cerebrospinal fluid lactate levels compared with age-matched nondemented individuals (Liguori *et al*., 2015). Thus, age-dependent changes in brain pH might be of pathophysiological significance in several neurodegenerative diseases, including HD (Jenkins *et al*., 1993; Madji Hounoum *et al*., 2017; Zheng *et al*., 2021). Importantly, changes in pH in the aging brain suggest that aging yeast and mammalian cells are subjected to similar environmental and cellular stresses. Thus, our data suggest that the yeast model can be an excellent model to decipher the mechanisms by which changes in extracellular pH can regulate proteotoxic stress in aging and human diseases.

## MATERIAL AND METHODS

### Drugs and Chemicals

Propidium iodide (P3566), acetic acid (9526), MES (J60763) were from Thermo.

### Strains and plasmid

Yeast strains and plasmids are listed in **Supplemental Table 1**.

### Culture conditions

Chronological aging experiments were carried out as previously described (Chadwick *et al*., 2016a, 2020c; Bari *et al*., 2023). Yeast cells expressing Htt^ex1^-CFP from the *GAL1* promoter were grown to saturation overnight in glucose (2%) synthetic minus histidine medium at 30°C in a rotating drum. The induction of the *GAL1* promoter was achieved by washing the saturated cells 3 times in galactose (2%) synthetic complete (SC) media lacking histidine followed by resuspending an aliquot of the washed cells in fresh media. All other cell types used were grown in glucose (2%) SC media with specific amino acid dropouts appropriate for the plasmid used. Cell viability assays were performed following the protocol as outlined by (Chadwick *et al*., 2016b). Cells were washed and then resuspended in phosphate-buffered saline (PBS) supplemented with 1 mM of propidium iodide. The positive control was prepared by boiling the cells for 10 min prior to resuspending them in the propidium iodide staining solution. Unstained cells were used as a negative control. All the samples were incubated in 96-well plates at room temperature for 10 min before being imaged on a Gel doc system (Bio-rad). The optical density (OD_600_) was assessed using a BioTek Epoch 2 microplate reader. Yeast cells expressing the protein of interest were cultured for several days and the chronologically aged cells viability was measured at different time intervals throughout the aging process. Alternatively, the CLS assay was conducted using medium buffered with 0.1 M MES buffer, or in high/low pH buffered media adjusted with NaOH. The acetic acid sensitivity was determined by culturing the cells in synthetic complete media for 4 days, and subsequently treated with acetic acid for 200 min at concentrations up to 0.08mM. Cell viability was then measured using propidium iodide. Survival rates were computed by measuring the mean gray value from images obtained along with OD_600_ using the ANALYSR program (Chadwick *et al*., 2016c). pH was measured using a Mettler-Toledo pH meter equipped with a micro pH electrode for small volume sampling.

### Growth assays

The OD_600_ of the overnight cell cultures were measured using a spectrophotometer. After adjusting the OD_600_ of the cell culture to 0.1, 5-fold serial dilutions were spotted on agar plates. Plates were incubated for 2 days at 30°C before taking images using a colony imager (S&P Robotics). The quantification was performed as previously described (Petropavlovskiy *et al*., 2020). Additionally, to assess the heat shock response, cells were also subjected to heat shock at 42°C for 1 hr before performing the growth assays.

### SDD-AGE

Experiments were carried as previously described (Jiang *et al*., 2017). Cells were harvested at days 1 and 3, with aliquots collected, centrifuged, and stored at −80°C. Cell lysis was performed using 100 μl and 200 μl of lysis buffer, respectively. The lysis buffer contained 100 mM Tris (pH 7.5), 200 mM NaCl, 1 mM EDTA, 5% glycerol, 1 mM DTT, 8 mM PMSF, and protease inhibitor cocktail. Each sample was combined with 100 mg of 0.5 mm acid-washed glass beads and mechanically lysed by vortexing, with 30-second cooling periods on ice between cycles. Protein concentrations were determined using Thermo Scientific Pierce BCA Protein Assay Kits. For gel electrophoresis, a 1.5% agarose gel was prepared in 1× TAE buffer, with 0.5% SDS added after complete dissolution. The running buffer consisted of 1× TAE containing 0.5% SDS. Samples were prepared using 4× sample buffer (2× TAE, 20% glycerol, 8% SDS, and bromophenol blue) diluted to 1× concentration, then boiled for 5 minutes at 100°C. Electrophoresis was conducted at 80 V for approximately 1.5 hours. Following electrophoresis, gels were rinsed with distilled water and then TBS buffer (20 mM Tris pH 7.5, 9 g NaCl per liter dH₂O). Proteins were transferred to PVDF membrane, which was subsequently blocked with BSA. The target protein was identified using antibody-antigen detection methodology.

### Fluorescence microscopy

Cells were spun down, washed and resuspended in PBS. Cells were diluted 10x and transferred to Nunc Lab-Tek chamber slides. Cells were imaged by utilizing a Zeiss Axiovert A1 wide-field fluorescence microscope with 63× 1.4 NA oil objective using a green fluorescent filter (489 nm excitation/508 nm emission) and an AxioCam ICm1 R1 CCD camera. ImageJ was used to analyze the images (Schneider *et al*., 2012)

### Flow cytometry

The mean fluorescent intensity of GFP/YFP reporters was determined using a BD FACS Celesta flow cytometer. A total of 50,000 cells were analyzed to determine the fluorescent intensities for each using a 488-nm blue laser. Mean fluorescence intensity was calculated using FlowJo.

### qRT-PCR

Total RNA was extracted using MasterPure Yeast RNA Purification Kit (Lucigen) as instructed in the manufacturer’s protocol. cDNA was synthesized by using the SuperScript IV VILO Master Mix (Thermo Fisher Scientific) as outlined in the manufacturer’s protocol. The primers used are listed in **Supplemental Table 2**. The relative expression of the gene of interest was quantified using the comparative Ct method and *TDH3* was used as a reference.

### RNA isolation and sequencing

Cells cultured both in normal media and in buffered media to saturation overnight, at 30°C in a rotating drum. Total RNA was extracted from cells using the RiboPure yeast kit (Thermo Fisher Scientific) as per the manufacturer’s manual. RNA samples were prepared with 3 replicates for each condition. Total RNA sequencing was carried out by Azenta Life Sciences. Stranded Illumina TruSeq cDNA libraries were generated from high-quality total RNA (RIN > 8) using poly dT. Libraries were sequenced on an Illumina HiSeq. The data have been deposited in NCBI’s Gene Expression Omnibus and are accessible through GEO Series accession number GSE297007 and GSE297008

### Quality control, trimming, read alignment, and differential gene expression analysis

The quality of the sequencing reads was assessed using FastQC (www.bioinformatics.babraham.ac.uk/projects/fastqc/). Next, Trimmomatic with its default settings were utilized to remove the low quality bases and adapter sequences (Bolger *et al*., 2014). The processed reads were aligned to the *S. cerevisiae* genome sequence (R64-2-1; www.yeastgenome.org/) with star (Dobin *et al*., 2013), ensuring uniquely mapped reads were only retained (Liao *et al*., 2014). The number of reads assigned for each gene were determined by featureCount. DESeq2 (Love *et al*., 2014) was used to analyze the differential gene expression analysis and genes with a Benjamini-Hochberg adjusted P-value of ≤0.05 were considered as significantly differentially expressed. GO analysis was performed using g:profiler (https://biit.cs.ut.ee/gprofiler/gost) (Reimand *et al*., 2007).

To identify Hsf1 and Msn2/4 targets, promoter analysis of differentially expressed genes (log2 fold change < 1 and *P*-value < 0.05) was performed using Yeastract (www.yeastract.com) (Monteiro *et al*., 2020).

### Intracellular pH measurements

Intracellular pH measurements using sfpHLuorin (addgene #115697) (Reifenrath and Boles, 2018) were done using flow cytometry using a BD FACS Celesta according to previously established methods (Triandafillou and Drummond, 2020). Briefly, cells expressing the sfpHLuorin were induced overnight in absence of methionine to induce expression of the reporter. To establish the calibration curve, cells were resuspended in a calibration buffer (50 mM NaCl, 50 mM KCl, 50 mM MES, 50 mM HEPES, 100 mM ammonium acetate, 10 mM 2-deoxyglucose; pH adjusted with HCl or KOH. 10 mM (1000x) nigericin in 95% EtOH was added just before buffer use to a final concentration of 10 μM.) at 0.5 pH unit intervals between pH 4.5 and pH 8.5. The median fluorescence ratio between the 405:525/50 channel (BV510, pHluorin 405) and the 488:525/50 channel (FITC, pHluorin 488) was calculated, and these data points were subsequently fitted to a sigmoidal function.

## ACKNOWLEDGEMENTS

PL is supported by an NSERC Discovery Grant (RGPIN-2022-05267), CIHR Project Grants (PJT 168882 and ARB 192062) and a Western CIHR Accelerator Grant. KAB was supported by an Ontario Graduate Scholarship. MLD is supported by an NSERC Discovery Grant (RGPIN-2024-05867). We thank Kevin Morano (UTHealth Houston) and David Pincus (University of Chicago) for yeast strains. Figure 1A was created using Biorender (Created in BioRender. Lajoie, P. (2025) https://BioRender.com/aiz9aec). We thank Donovan McDonald for useful discussions and comments on the manuscript.

---

**Supplemental Figure 1:**
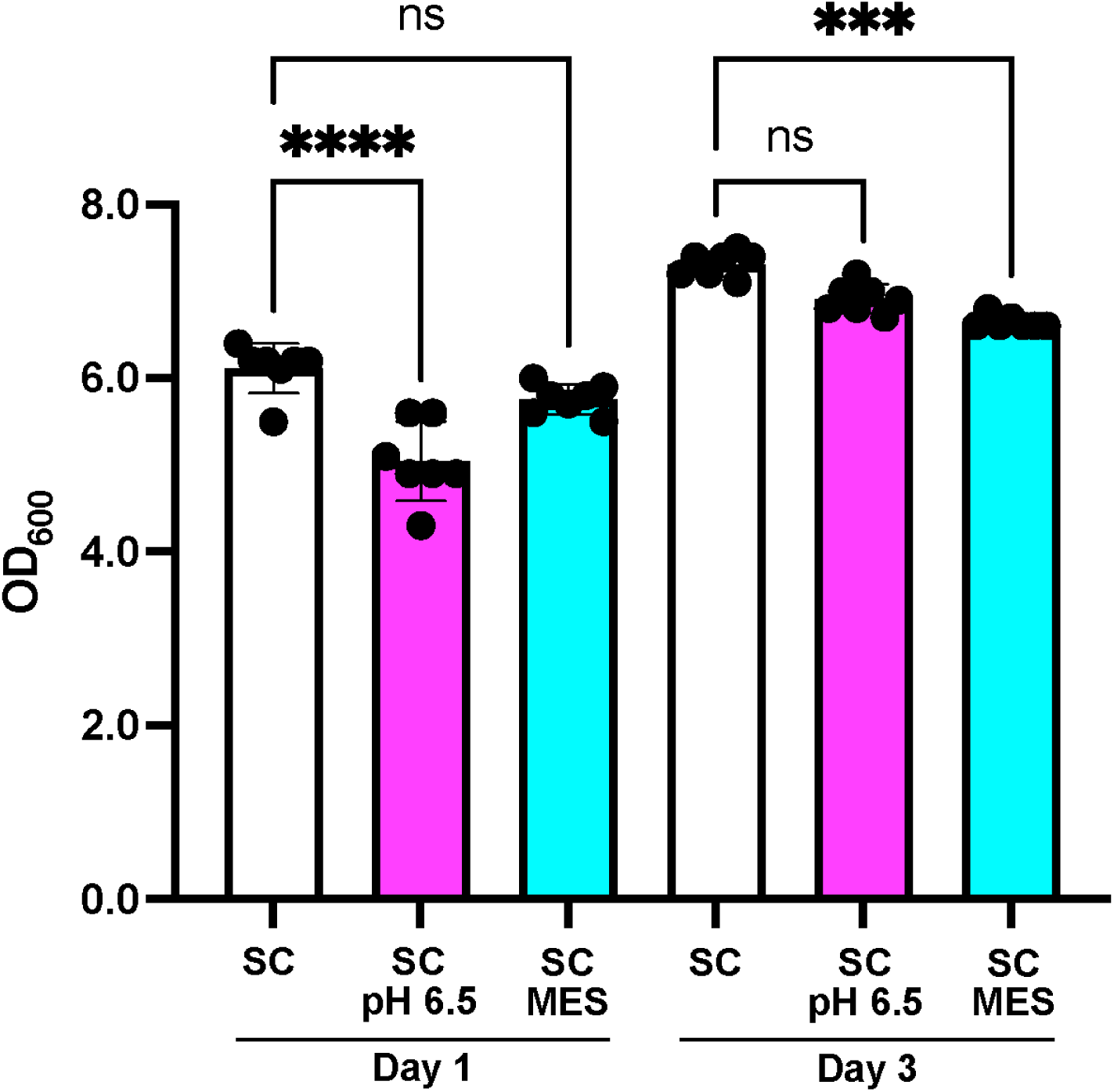
Growth in buffered media has minimal effect on cell density. Wild-Type cells were initially inoculated in regular SC media (pH 5.0), SC media buffered to pH 6.5 or media buffered with MES and grown for 1 or 3 days. Cell density (OD600) was measured using a spectrophotometer. (n= 7-10) +/-SEM). ***p<0.005 ****p<0.001. (Anova followed by Tukey’s multi comparison test).

**Supplemental Figure 2:**
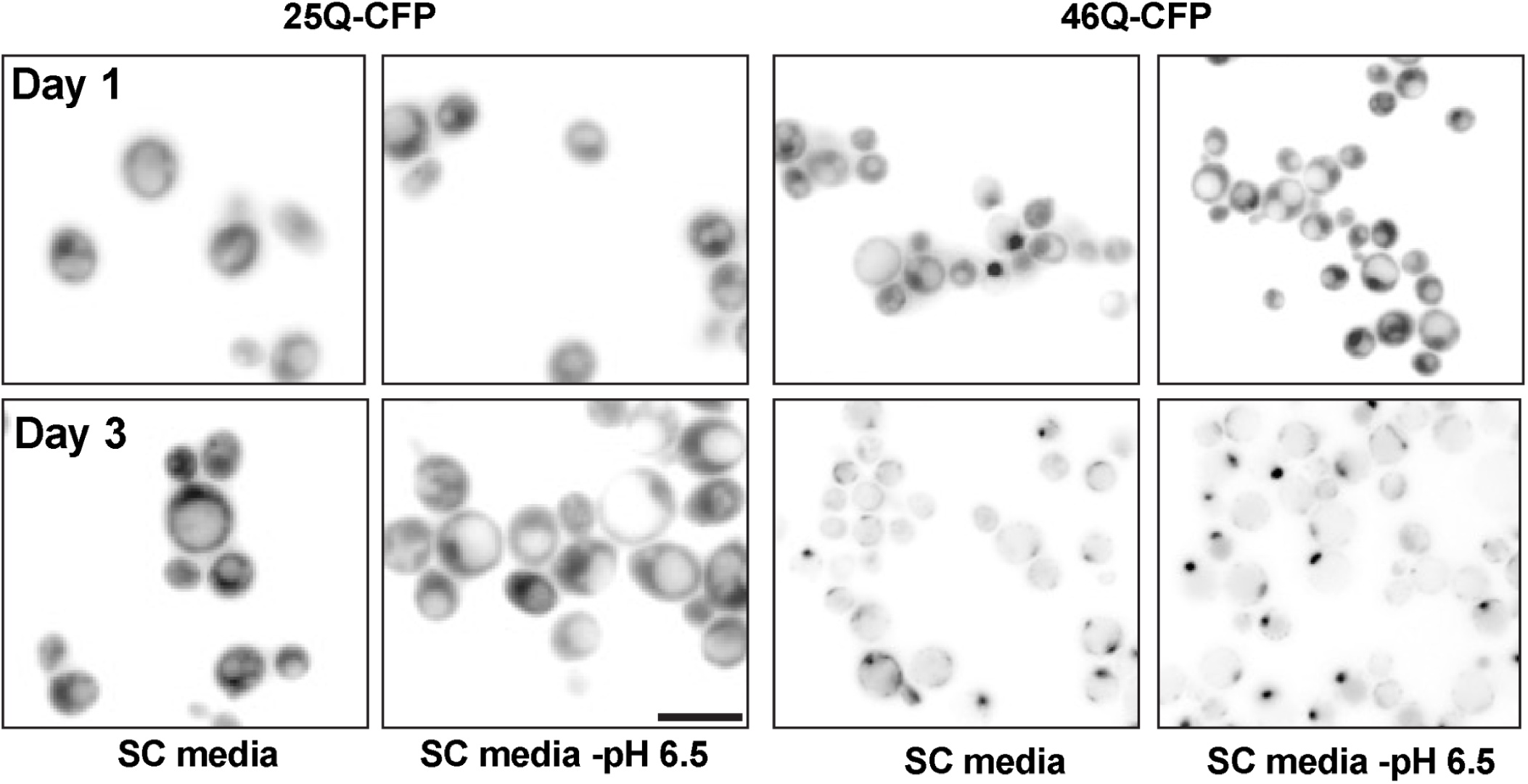
Increased presence of polyQ IBs in old cells cultured in media buffered to pH 6.5. Cells expressing 25Q-CFP or either expanded 46Q-CFP were processed for CLS in standard SC media or media buffered to pH 6.5 containing galactose. Fluorescence images were collected on day 1 and 3. Bar: 10µm

**Supplemental Figure 3:**
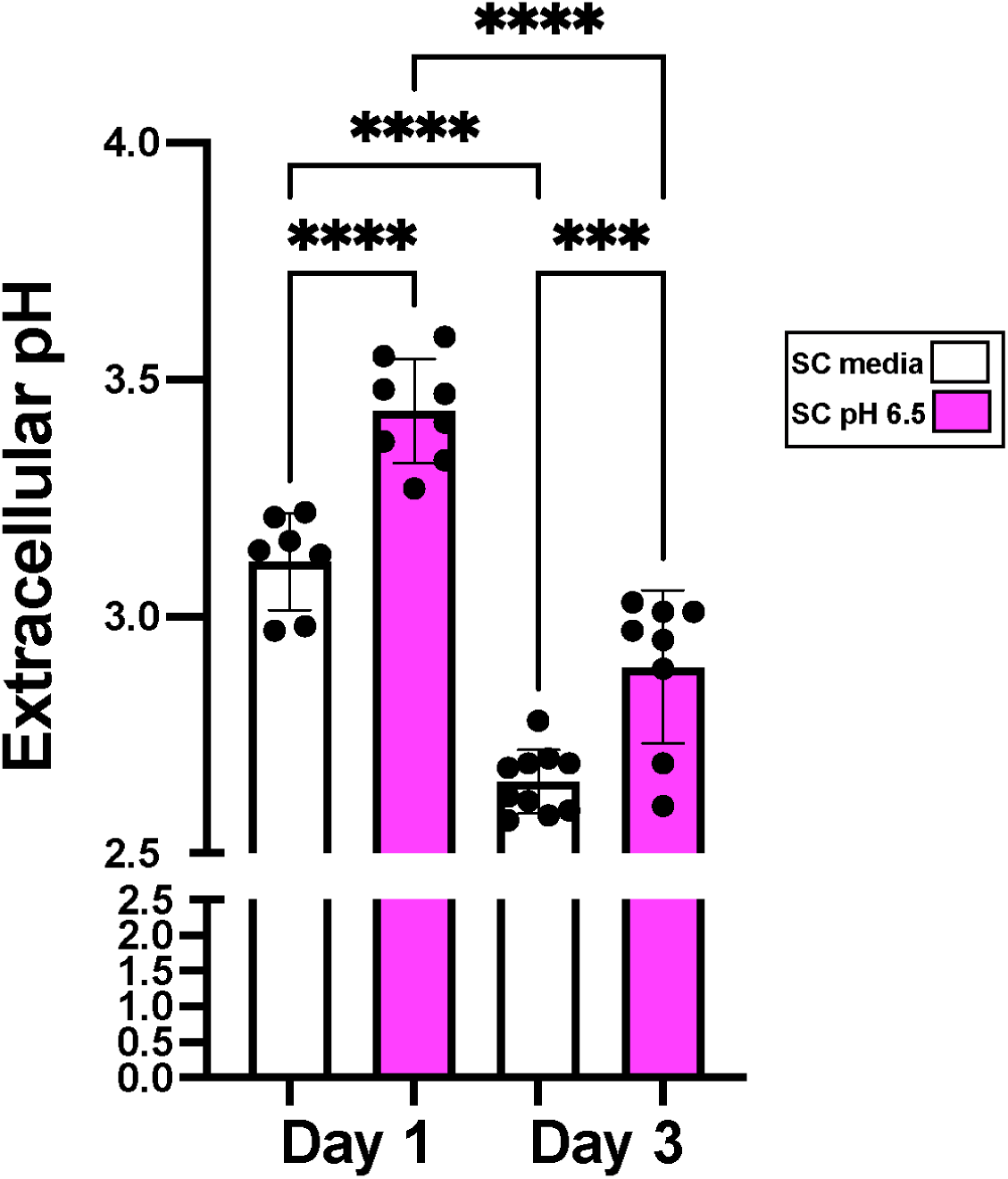
Extracellular pH still decreases during aging in media initially buffered to pH 6.5. Wild-Type cells were initially inoculated in regular SC media (pH 5.0) or SC media buffered to pH 6.5 and grown for 1 or 3 days and the extracellular pH was measured with a pH meter. (n= 7-10) +/-SEM). ***p<0.005 ****p<0.001. (Anova followed by Tukey’s multi comparison test).

**Supplemental Figure 4:**
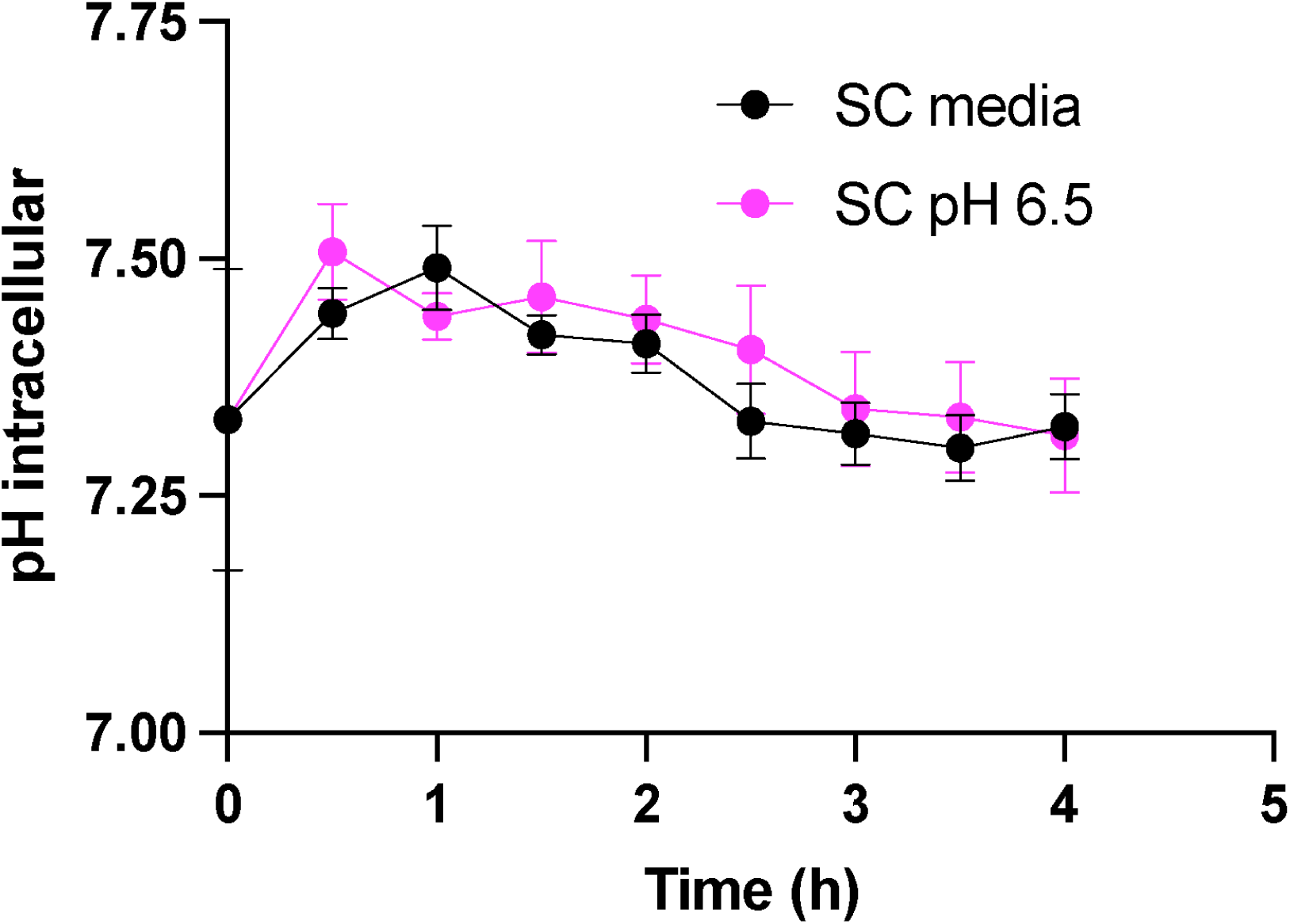
Buffering growth media to pH 6.5 does not affect intracellular pH. Wild-Type cells expressing sfpHLuorin were initially inoculated in regular SC media (pH 5.0) or SC media buffered to pH 6.5 and grown for 4 hours. Cytosolic pH was analysed every 30 min using flow cytometry.

**Supplemental Figure 5:**
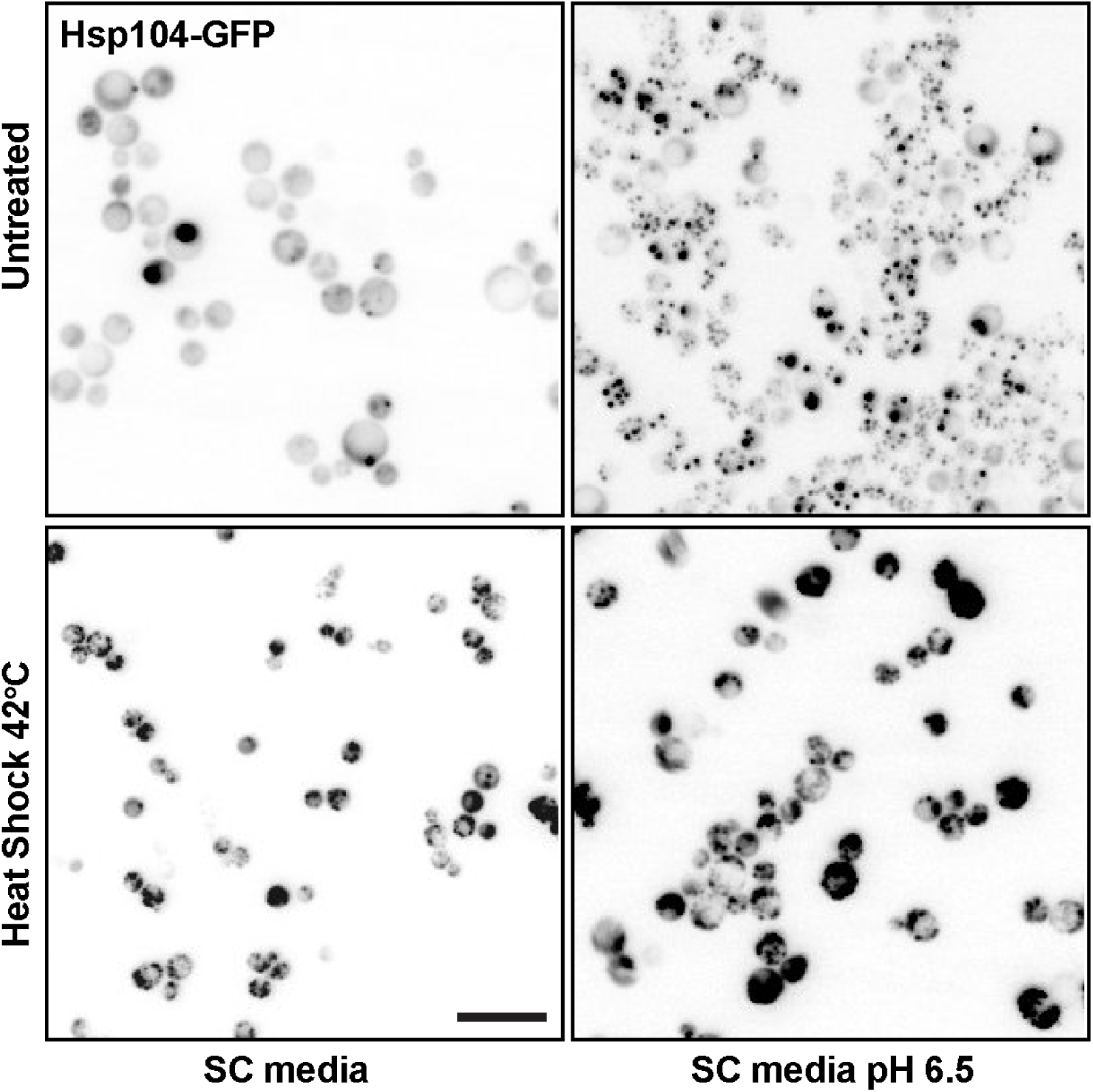
Growth at pH 6.5 induced aggregation of Hsp104. Cells expressing Hsp104-GFP were grown to early log phase in SC media or SC media buffered to pH6.5 and subsequently subjected to heat shock at 42°C for 1hr and fluorescence images were collected. Bar: 10µm

**Supplemental Figure 6:**
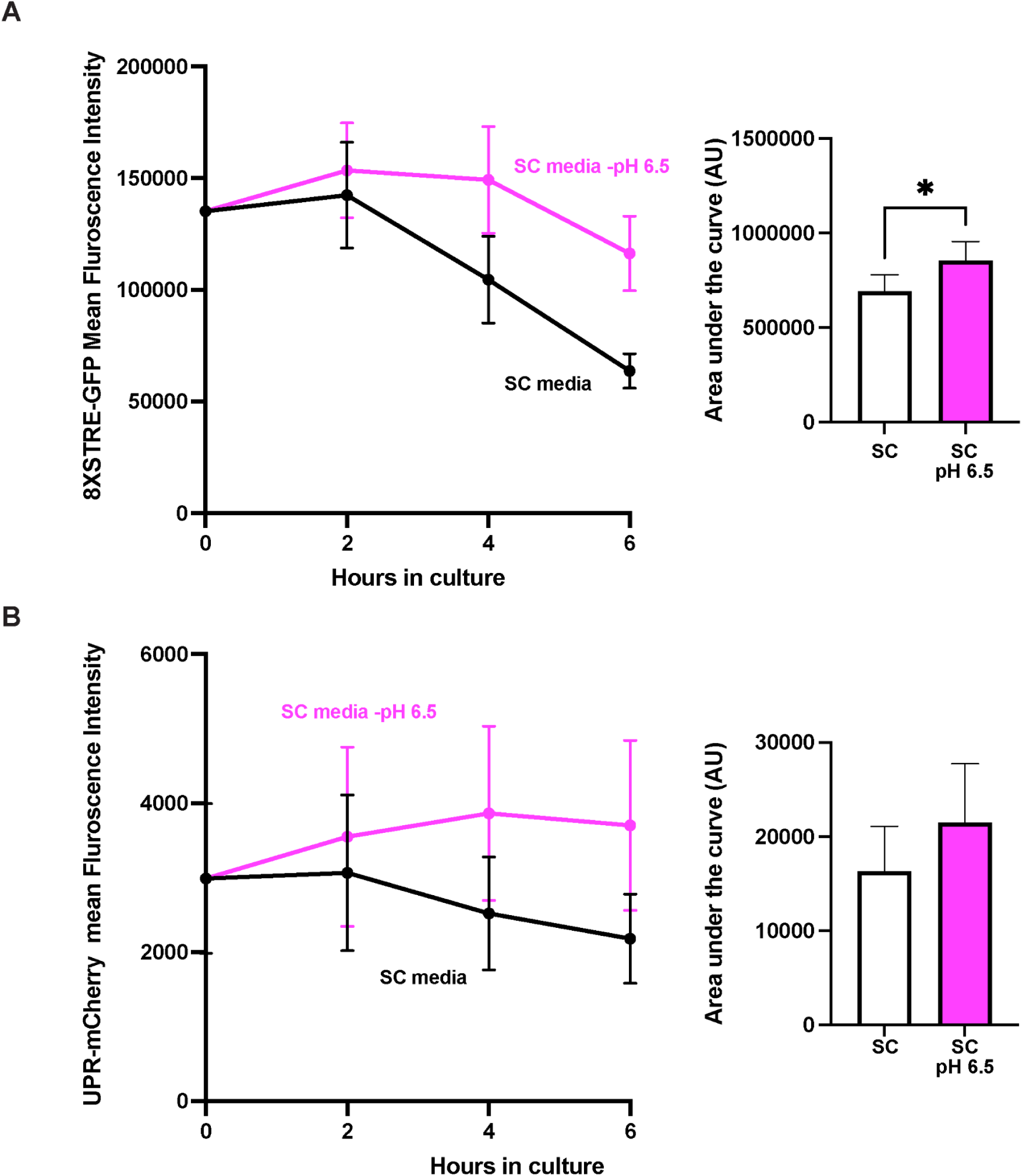
Growth at higher pH has modest effect on activation of proteotoxic stress responses other than the HSR. Fresh culture of cells expressing either a GSR responsive 8xSTRE-GFP reporter (**A**) or the UPR-mCherry (**B**) were grown overnight and transferred to either standard SC media or SC media buffered to pH 6.5 (Day 0) and analysed by flow cytometry at different time intervals. Mean fluorescence intensity of each reporter is shown in the graph (n=3). Area under the curves were calculated and presented in bar graphs. *p<0.05 (Student’s T-test).

**Supplemental Figure 7:**
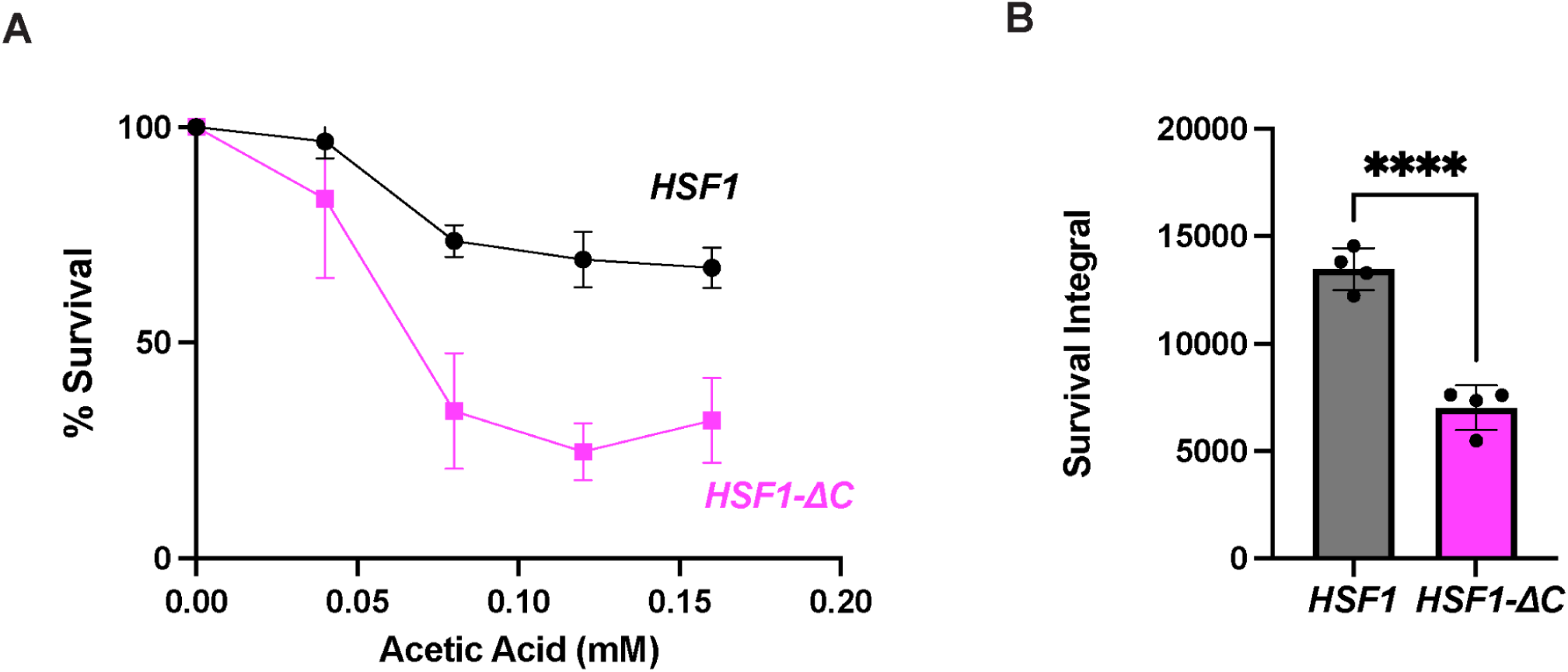
*HSF1ΔC* cells display increased sensitivity to acetic acid. Wild Type *HSF1* or *HSF1-ΔC* cells were grown for 2 days and treated with various concentrations of acetic acid. Viability was assessed with propidium iodide. **A)** Survival percentages normalized to untreated cells are shown on the graph (n=3 +/- SEM). **B)** The area under the curve (survival integral) is shown in the bar graph (n=3 +/- SEM). ****p<0.0001 (Anova followed by Tukey’s multi comparison test).

**Table S1:** Strains and plasmids used in this study.

| Strain Name | Genotype | Plasmid | Source |
| --- | --- | --- | --- |
| Wild-type W303a | *W303a derivative*<br>*MATa ade2-1 can1-100 trp1-1 leu2-3 his3-11 ura3-1* | - | (Thomas and Rothstein, 1989) |
| YPL016 | *W303a derivative*<br>*MATa ade2-1 can1-100 trp1-1 leu2-3 his3-11 ura3-1* | pRS303<br>*GAL1+FLAG-25QΔpro-CFP* | (Duennwald *et al.*, 2006b) |
| YPL017 | *W303a derivative*<br>*MATa ade2-1 can1-100 trp1-1 leu2-3 his3-11 ura3-1* | pRS303<br>*GAL1+FLAG-46QΔpro-CFP* |  |
| YPL018 | *W303a derivative*<br>*MATa ade2-1 can1-100 trp1-1 leu2-3 his3-11 ura3-1* | pRS303<br>*GAL1+FLAG-72QΔpro-CFP* |  |
| YPL104 | *W303a derivative*<br>*MATa ade2-1 can1-100 trp1-1 leu2-3 his3-11 ura3-1* | pRS424<br>*MET13+FLAG-25QΔpro-CFP* |  |
| YPL105 | *W303a derivative*<br>*MATa ade2-1 can1-100 trp1-1 leu2-3 his3-11 ura3-1* | pRS424<br>*MET13+FLAG-103QΔpro-CFP* |  |
| YPL807 | *W303a derivative*<br>*MATa ade2-1 can1-100 trp1-1 leu2-3 his3-11 ura3-1* | pRS426-*HSP104-GFP* | This study |
| YPL044 | *W303a derivative*<br>*MATa ade2-1 can1-100 trp1-1 leu2-3 his3-11 ura3-1* | pRS426-8XSTRE-GFP | This study |
| YPL033 | *W303a derivative*<br>*MATa ade2-1 can1-100 trp1-1 leu2-3 his3-11 ura3-1* | pRS316-UPR-RFP<br>Addgene #20132 | (Merksamer *et al.*, 2008) |
| DPY144 | *w303a; 4xHSE-Venus::LEU* | - | (Garde *et al.*, 2023) |
| YPL039a | *W303-1a hsf1::LEU2*<br>(*pRS314-HSF*) | pRS425<br>*MET13+FLAG-25QΔpro-CFP* | This study |
| YPL039b | *W303a hsf1::LEU2*<br>(*pRS314-HSF*) | pRS425<br>*MET13+FLAG-103QΔpro-CFP* |  |
| YPL040a | *W303a hsf1::LEU2*<br>[*pRS314-HSF(1-583)*] | pRS425<br>*MET13+FLAG-25QΔpro-CFP* |  |
| YPL040b | *W303a hsf1::LEU2*<br>[*pRS314-HSF(1-583)*] | pRS425<br>*MET13+FLAG-103QΔpro-CFP* |  |

**Table S2:**
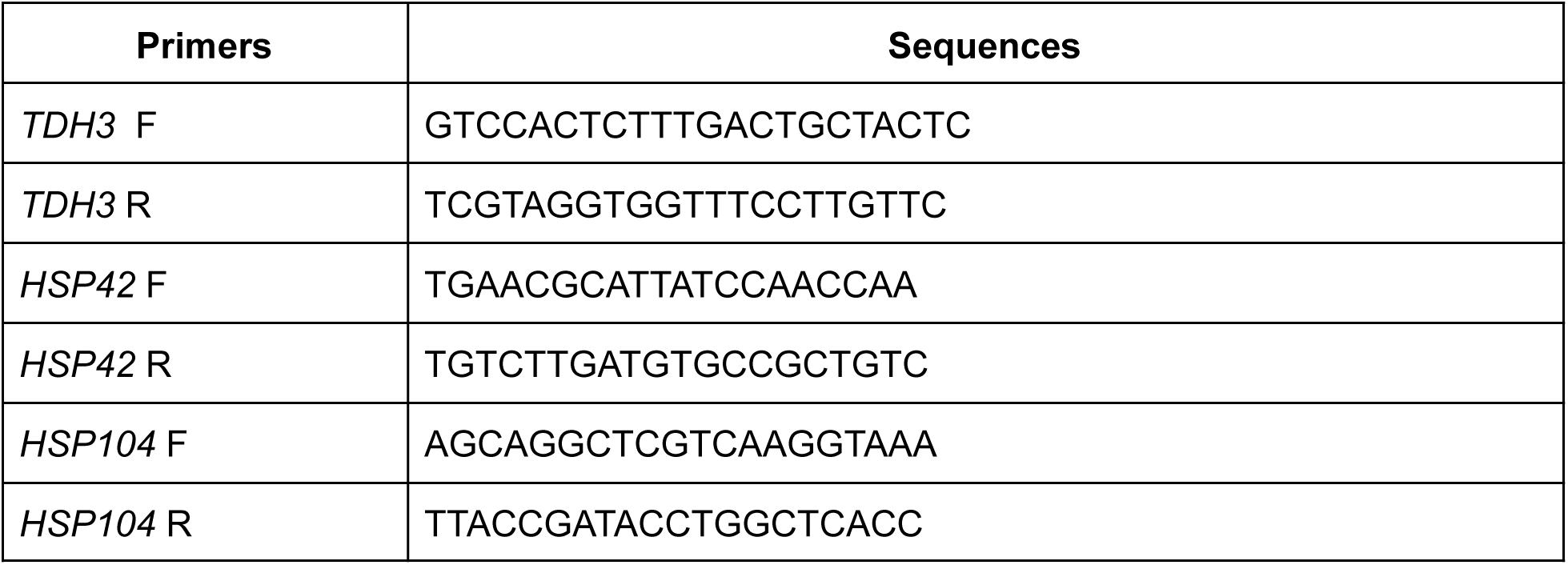
Primers used in this study.

